# Catalytic and non-catalytic mechanisms of histone H4 lysine 20 methyltransferase SUV420H1

**DOI:** 10.1101/2023.03.17.533220

**Authors:** Stephen Abini-Agbomson, Kristjan Gretarsson, Rochelle M. Shih, Laura Hsieh, Tracy Lou, Pablo De Ioannes, Nikita Vasilyev, Rachel Lee, Miao Wang, Matthew Simon, Jean-Paul Armache, Evgeny Nudler, Geeta Narlikar, Shixin Liu, Chao Lu, Karim-Jean Armache

## Abstract

The intricate regulation of chromatin plays a key role in controlling genome architecture and accessibility. Histone lysine methyltransferases regulate chromatin by catalyzing the methylation of specific histone residues but are also hypothesized to have equally important non-catalytic roles. SUV420H1 di- and tri-methylates histone H4 lysine 20 (H4K20me2/me3) and plays crucial roles in DNA replication, repair, and heterochromatin formation, and is dysregulated in several cancers. Many of these processes were linked to its catalytic activity. However, deletion and inhibition of SUV420H1 have shown distinct phenotypes suggesting the enzyme likely has uncharacterized non-catalytic activities. To characterize the catalytic and non-catalytic mechanisms SUV420H1 uses to modify chromatin, we determined cryo-EM structures of SUV420H1 complexes with nucleosomes containing histone H2A or its variant H2A.Z. Our structural, biochemical, biophysical, and cellular analyses reveal how both SUV420H1 recognizes its substrate and H2A.Z stimulates its activity, and show that SUV420H1 binding to nucleosomes causes a dramatic detachment of nucleosomal DNA from histone octamer. We hypothesize that this detachment increases DNA accessibility to large macromolecular complexes, a prerequisite for DNA replication and repair. We also show that SUV420H1 can promote chromatin condensates, another non-catalytic role that we speculate is needed for its heterochromatin functions. Together, our studies uncover and characterize the catalytic and non-catalytic mechanisms of SUV420H1, a key histone methyltransferase that plays an essential role in genomic stability.

## Introduction

Histone lysine methyltransferases play key roles in transcription, replication, and repair, and their dysregulation is found in diseases such as cancer (Husmann and Gozani, 2019; Martin and Zhang, 2005). Methylation of histone H4 at lysine 20 (H4K20me) is a highly abundant and conserved histone modification (Pesavento et al., 2008; Schotta et al., 2004). The enzyme paralogs, SUV420H1 and SUV420H2, catalyze the di- and trimethylation of H4K20 (H4K20me2 and me3) and require the H4K20me1 deposited by Set8 monomethyltransferase (Nishioka et al., 2002; Schotta et al., 2004; Schotta et al., 2008). Whereas SUV420H1 is highly expressed, and its deletion results in perinatal lethality, SUV420H2 plays more peripheral roles (Schotta et al., 2008). SUV420H1 is dysregulated in neurodevelopmental and neuropsychiatric disorders, and its mutations, deletions, and amplifications are found in a host of cancers (Brohm et al., 2019; Paulsen et al., 2022; Wang et al., 2021). SUV420H1 plays critical roles in proliferation, cell cycle progression, heterochromatin formation, DNA damage response, and telomere elongation, suggesting important roles in regulating chromatin structure and function (Jorgensen et al., 2013).

SUV420H1 is targeted to heterochromatin through interactions with heterochromatin protein 1 (HP1) (Schotta et al., 2004). Together with SUV420H2, it establishes H4K20me3, which plays a poorly understood role in heterochromatin silencing. SUV420H1-catalyzed H4K20me2 also plays a role in DNA damage and p53 signaling during double-strand break repair by recruiting 53BP1, which binds the H4K20me2 mark using its Tudor domain (Paquin and Howlett, 2018; Wilson et al., 2016). SUV420H1 also plays a vital function at the origins of replication, where it is recruited and stimulated by histone variant H2A.Z and deposits H4K20me2, a modification recognized by the BAH domain of Orc1 (Kuo et al., 2012; Long et al., 2020). The mechanism by which H2A.Z stimulates SUV420H1 activity is not well understood.

Besides catalyzing deposition of histone modifications, many histone-modifying enzymes possess non-catalytic functions (Illingworth et al., 2015; Lee et al., 2018; Sze et al., 2017). This leads to uncertainty regarding instructive versus correlative functions of histone modifications. However, non-catalytic mechanisms of histone lysine methyltransferases are poorly understood, precluding designing inhibitors and therapeutic strategies to target their dysregulation in disease. Whereas SUV420H1 and H2 double knockout mouse embryonic fibroblasts are hypersensitive to DNA damage, cells subjected to a SUV420 catalytic inhibitor are only mildly sensitive to DNA damaging agents (Bromberg et al., 2017; Schotta et al., 2008). This suggests a structural, non-catalytic role of SUV420 proteins. To understand how SUV420H1 dysregulation causes disease, it is critical to understand both the catalytic and non-catalytic activities of this enzyme on chromatin. We determined cryo-EM structures of SUV420H1 complexes with nucleosomes containing catalytic histone H2A or its variant H2A.Z. These structures reveal the molecular interactions involved in H4K20 methylation, and together with biochemical, biophysical, and cellular studies, uncover the mechanisms by which SUV420H1 regulates chromatin. Specifically, the discovery and molecular characterization of non-catalytic mechanisms of SUV420H1 are critical for understanding its effects on chromatin. Our direct visualization of how SUV420H1 affects nucleosome distortion and chromatin condensation opens new avenues to address its dysfunction in disease.

## Results

### SUV420H1 is stimulated by histone variant H2A.Z

To understand the mechanism by which histone variant H2A.Z regulates SUV420H1, we measured the activity of the catalytic fragment of SUV420H1 (residues 1-393, which also corresponds to full-length isoform 2) (Tsang et al., 2010) using nucleosomes reconstituted with a methyl-lysine analog of monomethylated histone H4 lysine 20 (H4K_C_20me1) and either histone H2A or histone variant H2A.Z. Endpoint methyltransferase assays revealed more methyltransferase activity on H2A.Z/H4K_C_20me1 than on nucleosomes containing H2A/H4K_C_20me1 (Figure 1A). Western blot analysis also showed more H4K_C_20me2 on H2A.Z nucleosomes than on H2A nucleosome substrates (Figure S1A). To further understand the mechanism, we performed enzyme kinetics using H2A/H4K_C_20me1 and H2A.Z/H4K_C_20me1 containing nucleosomes. We observed that H2A.Z containing nucleosomes result in a higher apparent unimolecular rate constant (*k*_cat_) compared with that of H2A nucleosomes, whereas the Michaelis constant (*K*_M_) for methyl transfer is similar for both substrates suggesting an allosteric stimulation of H4K20 methylation on H2A.Z nucleosomes (Figure 1B). Similar stimulation was observed using nucleosomes devoid of H4K_C_20me1 (Figures S1B and S1C). To understand which residues in H2A.Z are required for SUV420H1 stimulation, we focused on the docking domain of H2A.Z, which contains residues that contribute to the nucleosome acidic patch (Figure S1D) (Suto et al., 2000). The negatively charged acidic patch formed in part by residues contributed by H2A/H2A.Z emerged as a prominent interacting surface for chromatin-binding proteins (Armache et al., 2011; McGinty and Tan, 2015). Sequence substitution in H2A.Z introduces an aspartic acid (D94), increasing the charged surface of the acidic patch in H2A.Z-containing nucleosomes. To test the importance of the acidic patch in general and to understand how H2A.Z-specific residues impact the activity of SUV420H1, we performed activity measurements on either wild-type H2A or chimeric H2A that harbors H2A.Z-specific residues (N94D/K95S). We observed a significantly higher methylation activity of SUV420H1 on chimeric H2A nucleosome than on the wild-type H2A nucleosome, suggesting that the extended H2A.Z acidic patch stimulates SUV420H1 activity (Figure S1E).

**Figure 1.**
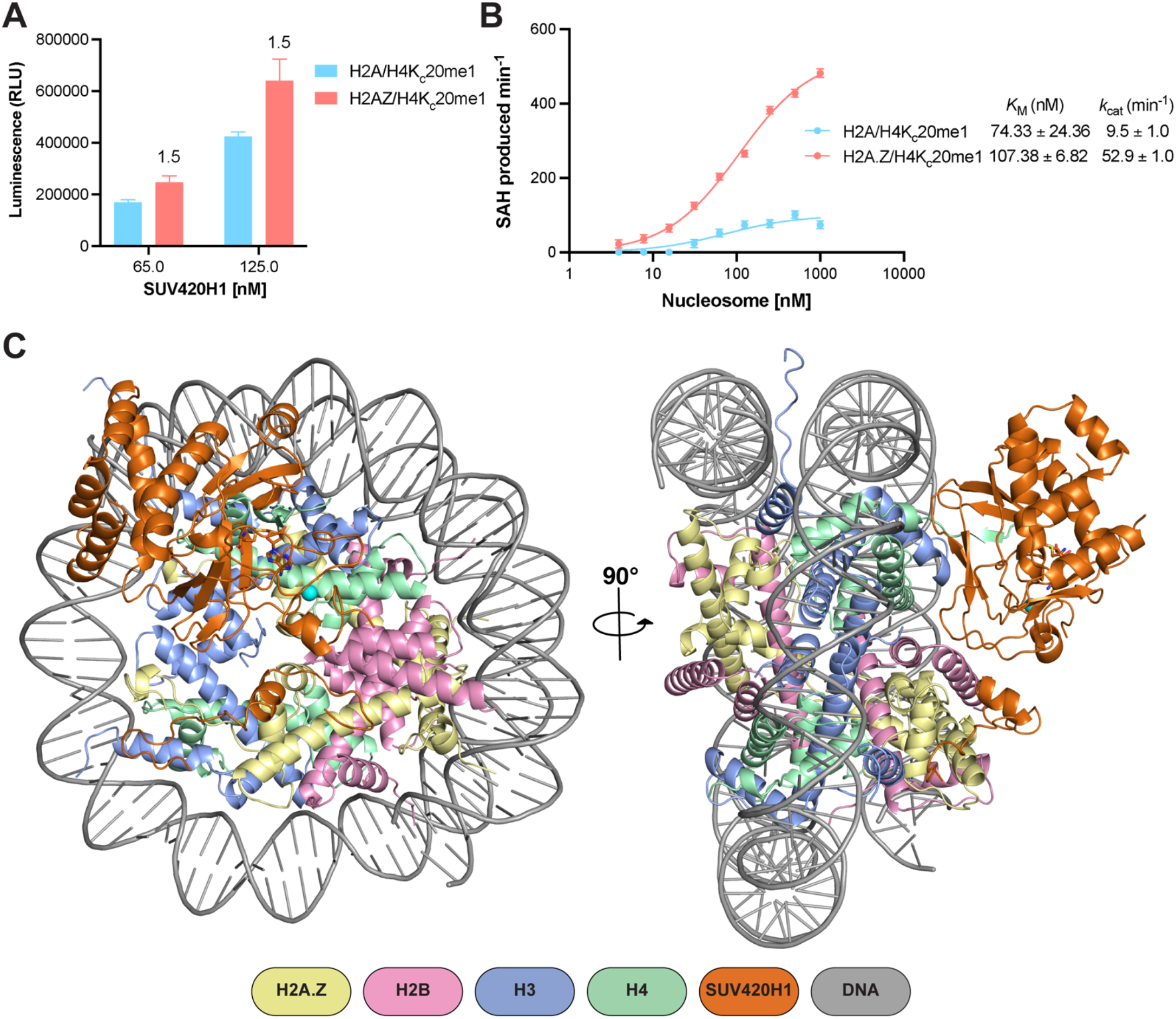
Activity and structure of SUV420H1-nucleosome complex. A. Catalytic activity of SUV420H1 on H2A and H2A.Z H4K_C_20me1 nucleosomes measured using an endpoint methyltransferase assay. Values above the bars represent the ratios between SUV420H1 activity measured on H2A.Z and H2A nucleosomes. B. Michaelis–Menten titrations of H4K_C_20me1 nucleosomes containing H2A and H2A.Z with SUV420H1. The *K*_M_ and *k*_cat_ values of the fitted data are reported in the graph. Each data point represents the mean ± SD from experiments repeated at least three times. C. Structures of SUV420H1-H2A.Z nucleosome complex shown in two different views. The color scheme for complex composition is indicated below the model.

### SUV420H1-H2A.Z nucleosome structure determination

To gain a mechanistic understanding of chromatin recognition and modification by SUV420H1, we assembled a complex of SUV420H1 (residues 1-393) and H2A.Z nucleosome. The nucleosomes contained a SET domain–stabilizing mutation H4K20M (Grau et al., 2021; Lewis et al., 2013; Shan et al., 2016; Valencia-Sanchez et al., 2021). Gel-filtration analysis confirmed the formation of the SUV420H1-H2A.Z nucleosome complex (Figure S2A). The sample was crosslinked using glutaraldehyde cross-linking gradient fixation (GraFix) (Stark, 2010) (Figure S2B), and cryo-EM data was collected on a 300kV Titan Krios microscope. We determined the structure of the SUV420H1-H2A.Z nucleosome complex to an overall resolution of 2.6 Å (Figures 1C, S3 and S4). We were able to unambiguously fit the crystal structure of the catalytic domain of SUV420H1 (residues 70–336) into the cryo-EM map (Figure S5) (Wu et al., 2013). The overall structure of the catalytic domain was highly similar to the previously determined SUV420H1 and SUV420H2 catalytic domains (Figures S6A and S6B) (Southall et al., 2014; Wu et al., 2013). The cryo-EM map did not show density for the unstructured N-terminus of SUV420H1 (residues 1–72); however, we observed the C-terminal density extending beyond the catalytic domain structures (residues 346–378) (Figure S5). Residues of the SUV420H1 not present in the crystal structure were modeled *de novo*, guided by the cryo-EM density and cross-linking mass spectrometry (XLMS) (Table S1) (O’Reilly and Rappsilber, 2018). Most of the crystal structure of SUV420H1 corresponded well with our cryo-EM map, except the N-terminal helical domain, which was rigid-body fitted because of low resolution in that region (Figure S5).

### Recognition of DNA and histone H4 by SUV420H1

SUV420H1 makes extensive intermolecular contacts with the nucleosome (Figure 2A). The enzyme engages the H4 tail via two loops, the ‘DNA-interacting loop’ and the ‘H4-interacting loop’ (Figure 2B). R286, within the DNA-interacting loop, forms a hydrogen bond with the phosphate backbone of DNA (Figures 2B and S7A). Consequently, an R286A mutation reduced the catalytic activity of SUV420H1, highlighting the importance of DNA interaction in the methylation of H4K20 (Figure 2B). H4-interacting loop residues S255, T256, and R257, establish multiple contacts with H4 residues D24 and N25 (Figure S7B). Mutations in this region of SUV420H1 are associated with decreased methyltransferase activity and are found in cancers (Brohm et al., 2019). The catalogue of somatic mutations in cancer (COSMIC) database reveals that S255F and K258E are frequently observed in breast invasive lobular carcinoma and uterine endometrioid carcinoma, respectively (Cerami et al., 2012). Our structure reveals that residue S255 makes sidechain interaction with D24 of histone H4 (Figure S7B) and K258 residue of SUV420H1 forms intramolecular interactions with the DNA-interacting loop (Figure S7C), perhaps contributing to its stabilization. The cryo-EM map allows unequivocal assignment of critical interactions between the H4 tail and SUV420H1 (Figure S7D). Hydrogen bonds and salt bridges mediate interactions between histone H4 tail and SUV420H1’s SET domain. The H4 tail in the SET domain presents a similar architecture to the previously published SUV420H2-H4 peptide structure (PDB: 4AU7) (Figure S6C) (Southall et al., 2014). In our structure, the H4 tail forms a parallel β sheet configuration with β4 of the SUV420H1 SET domain (Figure S6D). Histone H4 residues L22, V21, M20, and H18 form main chain contacts with SUV420H1 (Figure 2C). The hydrophobic side chain of H4K20M is pointed towards the SUV420H1 catalytic residue S251 and is close (∼5.2Å) to the methyl donor of the ligand S-adenosylmethionine (SAM) (Figures 2C and S6D). In agreement with other studies, H4 residues R19 and R17 form key sidechain interactions with SUV420H1 residues E239 and E314, respectively, contributing to substrate specificity (Figure 2C) (Kudithipudi et al., 2012; Southall et al., 2014). Consistent with our hypothesis that SUV420H1 E314 might be essential for the proper conformation of the H4 tail, a charge swap mutation E314K decreased the catalytic activity of SUV420H1 on both H2A and H2A.Z nucleosomes (Figure 2C). Our structure thus reveals the full extent of interactions between SUV420H1 with histone H4, and their roles in methylation activity.

**Figure 2.**
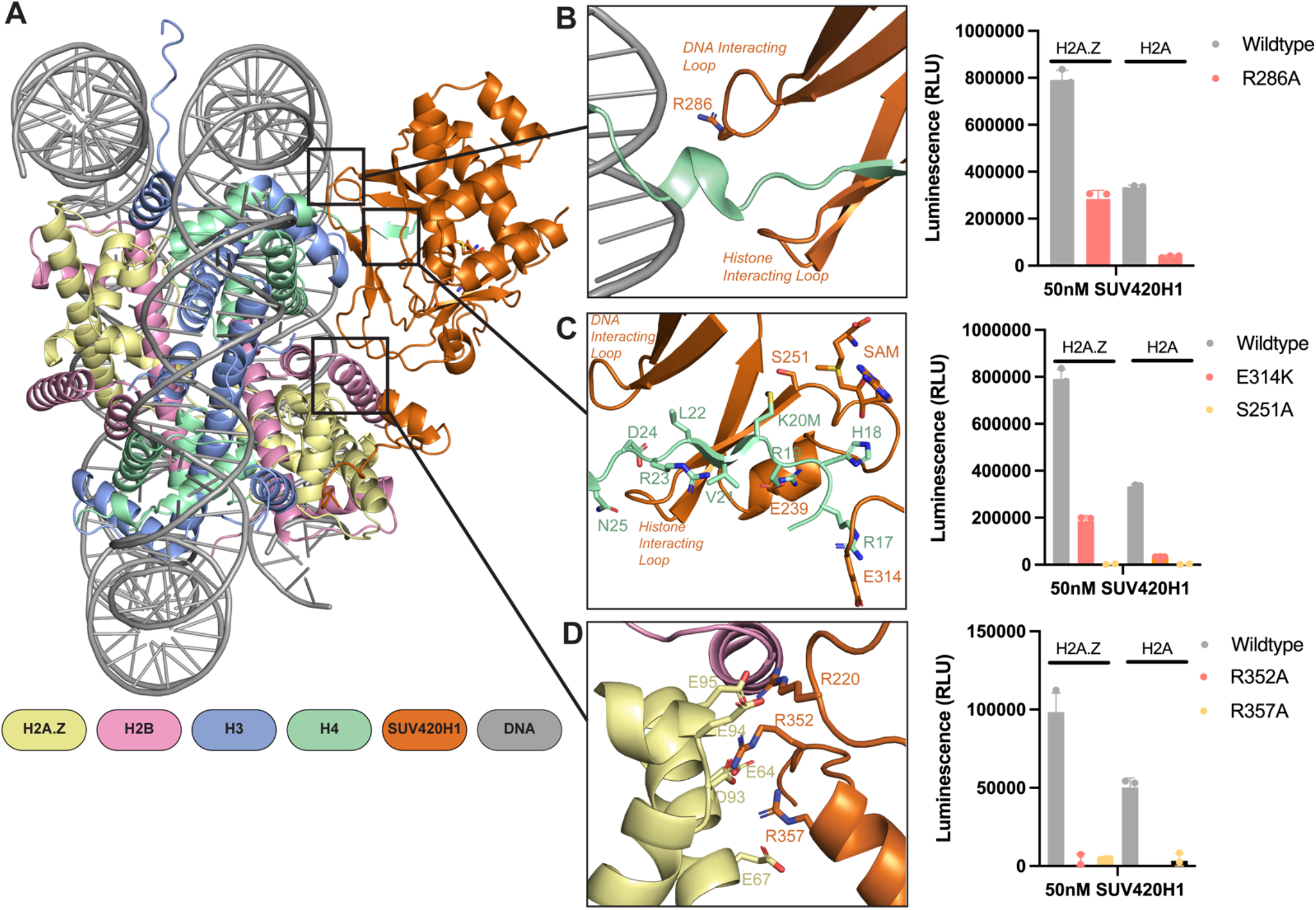
Interactions of SUV420H1 with H2A.Z nucleosome. A. Overview of the contacts between SUV420H1 and H2A.Z nucleosome. B. Close-up view of interactions between SUV420H1, histone H4, and DNA. On the right is a representative methyltransferase activity of wildtype SUV420H1 and R286A mutant on H2A and H2A.Z nucleosome. C. Detailed interactions of histone H4 with the SET domain of SUV420H1 with representative methyltransferase activity of wildtype SUV420H1 and mutants (E314K and S251A). D. Detailed interaction between SUV420H1 and acidic patch residues of H2A.Z nucleosome with representative methyltransferase activity showing the effect of mutants (R352A and R357A). For each methyltransferase assay, data points and error bars represent the mean ± SD from three independent experiments.

### Role of acidic patch in H4K20 methylation

Our cryo-EM map of SUV420H1 bound to H2A.Z nucleosomes presented density associated with the acidic patch interaction (Figure S5). We modeled the C-terminal region of SUV420H1 (residues 346–378), aided by cross-linking mass spectrometry (XLMS) (Table S1). In general, the acidic patch interacting region of SUV420H1 remains unstructured except for a short helix (residues 355–361) (Figure 2D). We observed a canonical acidic patch interaction involving R352 and R357 of SUV420H1 (Figure S8A). Both residues form a salt bridge with the H2A.Z residue E64, and interact with D93, E95, and E67 (Figure 2D). The functional importance of these SUV420H1 residues in H4K20 methylation is highlighted by the considerable loss of activity of their mutations (Figure 2D). On both edges of the acidic patch, SUV420H1 makes potential hydrogen bond interactions with H2A.Z and H2B (Figures S8B and S8C). Beyond the acidic patch interacting region, SUV420H1 R360 forms potential hydrogen bonds with N71 of H2A.Z, a residue conserved in H2A. An R360A mutation in SUV420H1 decreases enzymatic activity (Figure S8C). Thus, SUV420H1 stable anchoring to nucleosome by H2A/H2A.Z acidic patch and other proximal residues is crucial in the methylation of H4K20. Our cryo-EM model does not reveal direct contacts between SUV420H1 and H2A.Z-specific acidic patch residue (D97). We have observed that the loop between ß2 and ß3 in the SUV420H1-H2A.Z-nucleosome structure is more ordered compared to the SUV420H1-H2A nucleosome structure. (Figures S9A and 9B). Furthermore, R220, located in this loop, projects its side chain towards the acidic patch in the SUV420H1-H2A.Z structure (Figures 2D and S9C). Mutations and truncation at R220 are observed in uterine endometrioid carcinoma, pancreatic adenocarcinoma, and cutaneous melanoma (Cerami et al., 2012). An R220A mutation reduces the activity of the protein on both H2A.Z as well as H2A nucleosomes (Figure S9D). These observations suggest that the presence of H2A.Z, which contains an extended acidic patch, may enhance the stability of the enzyme-substrate complex and possibly lead to a higher catalytic activity.

### SUV420H1 induces detachment of terminal nucleosomal DNA from the histone octamer

The cryo-EM map of the SUV420H1-H2A.Z nucleosome complex showed terminal nucleosomal DNA detached from the octamer (Figure 3). The detachment of terminal DNA is correlated with the presence of SUV420H1 (Figures 3A and 3B). Two major cryo-EM map reconstructions of SUV420H1-nucleosome complexes were obtained; the first had one SUV420H1 bound on a single face of the nucleosome and showed the detachment of DNA at SHL +5, SHL +6, and SHL +7 (Figure 3A). The second showed SUV420H1 bound to both faces of the nucleosome with a detachment of DNA at both entry and exit (SHL ±5, SHL ±6, and SHL ±7) (Figure 3B). A similar detachment was observed in the structure of SUV420H1 bound to H2A nucleosome but not in the structures of H2A.Z or H2A nucleosomes alone, supporting the SUV420H1-mediated nucleosomal DNA detachment (Figures 3C, S10, S11A and S11B). Thus, the binding of SUV420H1 to nucleosome facilitates DNA detachment (Figure 3D). In a 3D classification of cryo-EM data, we can observe an unexpected density on the nucleosome surface close to αN helix of histone H3 in the subset(s) where terminal DNA is detached from the histone octamer (Movie1). The additional density can be attributed to the C-terminal residues (362–393) of SUV420H1 (Figures 4A and 4B). We used crosslinking mass-spectrometry analyses to corroborate that these residues in SUV420H1 interact with histone H3 (Figure 4C and Table S1). The cryo-EM map resolution limits our ability to determine sidechain interactions between the C-terminal region of SUV420H1 and the nucleosome. Its position is in proximity to the αN helix of histone H3, which harbors K56, modification of which was shown to influence the unwrapping dynamics of nucleosomes (Zhou et al., 2019). It is noteworthy that the C-terminal region of SUV420H1 contains several charged and polar residues, which could interact with H3 αN helix to weaken histone–DNA contacts. To examine whether the SUV420H1 C-terminus was essential for enzymatic activity, we performed endpoint methyltransferase assays and kinetic analyses to compare wildtype and ΔC-terminus mutant SUV420H1 (residues deleted 362–393) and observed only a minimal change in activity (Figures S12A and S12B). To determine whether the C-terminus of SUV420H1 is sufficient for DNA detachment, we pursued structural studies of a peptide containing this region bound to nucleosomes. We used a peptide containing the acidic patch binding residues and histone H3–interacting residues (N346–S378) (Figure S12C), reconstituted the complex with H2A.Z-nucleosomes, and determined the structure of the peptide-bound nucleosome. The 3 Å cryo-EM map shows detachment of DNA from the nucleosome at the entry site (Figure 4D). Further processing of the cryo-EM data revealed the fully wrapped and unwrapped particles but 3D variability analysis shows that a 3D class with detached DNA correlated with increased SUV420H1 C-terminal peptide occupancy close to H3 αN helix (Movie 2). These results collectively suggest that SUV420H1 C-terminal interactions with the histone H3 αN helix promote DNA detachment without affecting H4K20 methylation, indicating that we discovered a non-catalytic activity of SUV420H1.

**Figure 3.**
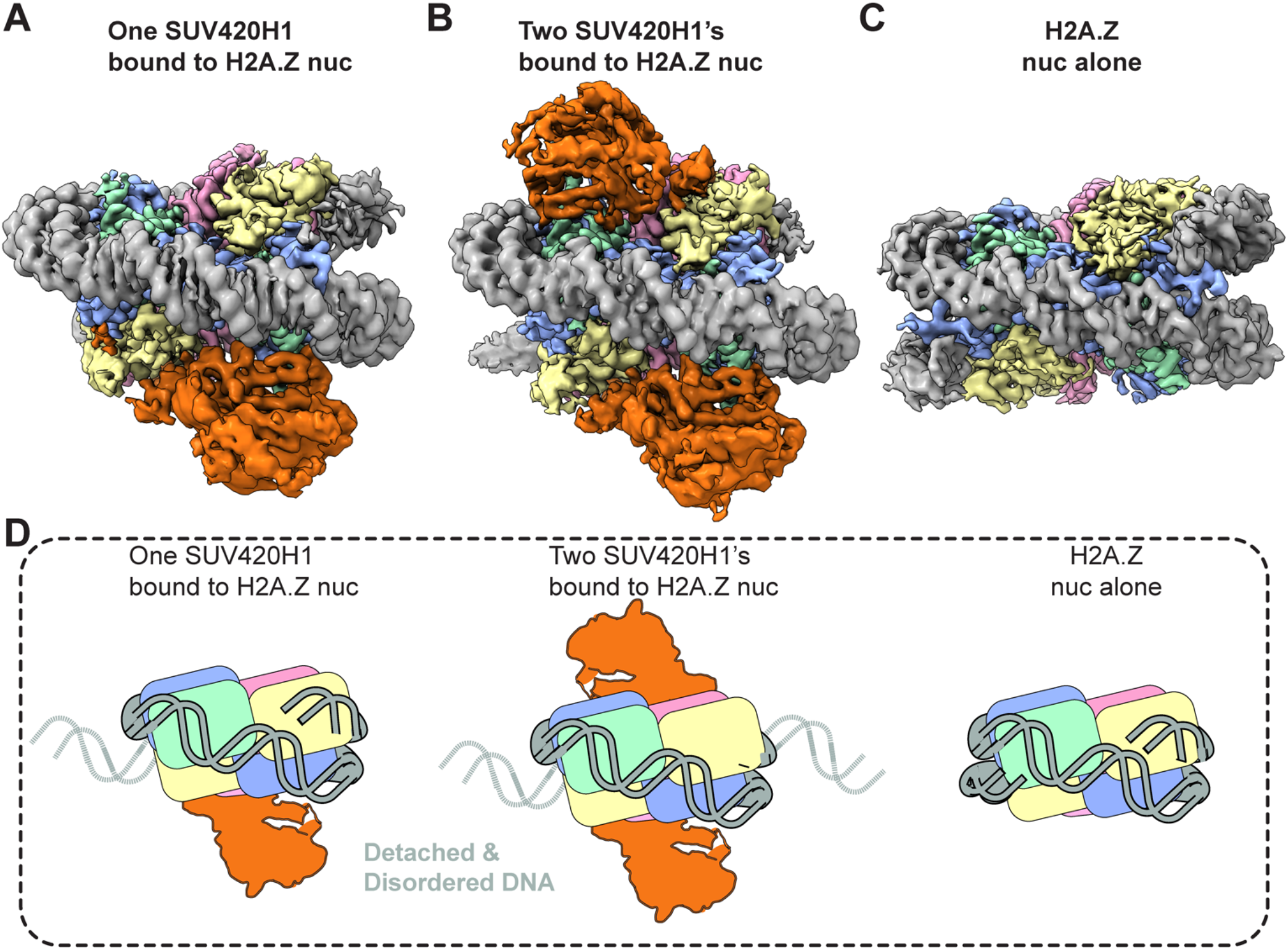
Distinct cryo-EM reconstructions of SUV420H1 bound to H2A.Z nucleosome. A. SUV420H1 bound to one face of the H2A.Z nucleosome. B. SUV420H1 bound to both faces of the H2A.Z nucleosome. C. H2A.Z nucleosome alone. D. Cartoon depiction showing how detachment of DNA correlates with SUV420H1 binding occupancy.

**Figure 4.**
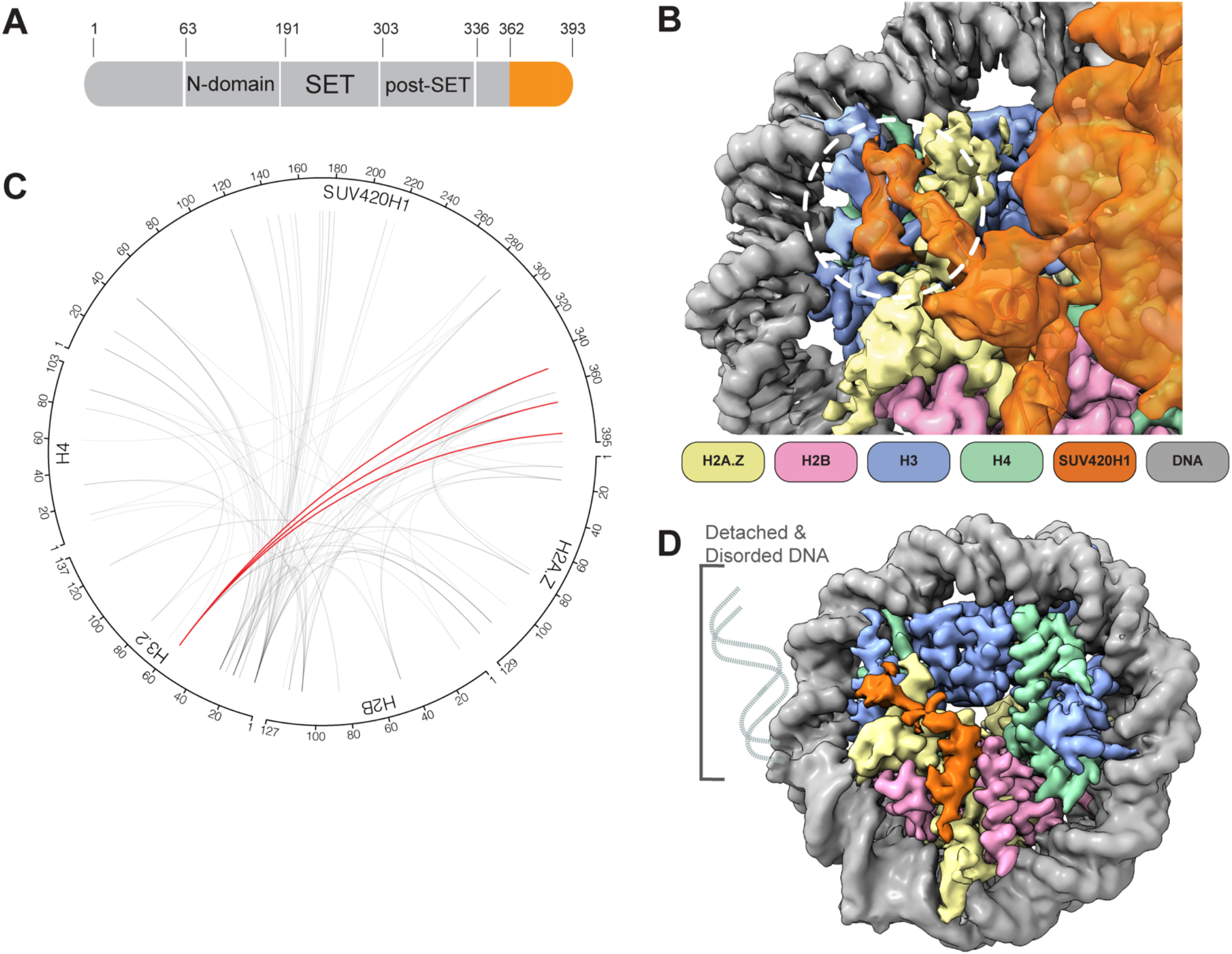
SUV420H1 C-terminus and nucleosome interactions. A. Diagram of SUV420H1 highlighting the C-terminal region colored in orange. B. Close-up view showing the cryo-EM density of SUV420H1 C-terminus, histones H3 and H2A.Z. The interacting region between SUV420H1 C-terminus and αN helix of histone H3 is circled in white. C. Circular diagram showing intermolecular crosslinks found in the SUV420H1-H2A.Z nucleosome complex. The crosslinks between SUV420H1 C-terminus and αN helix of histone H3 are shown in red, and all other crosslinks are shown in grey lines. The intramolecular crosslinks are not presented for simplicity. D. Cryo-EM reconstruction of SUV420H1 C-terminal peptide bound to H2A.Z nucleosome, showing DNA detachment.

### SUV420H1 remodels and condenses chromatin arrays

Our cryo-EM results suggest that SUV420H1 binds to the nucleosome and alters its structure independent of the enzymatic activity of SUV420H1. We employed single-molecule force spectroscopy to obtain the “mechanical fingerprint” of SUV420H1-bound nucleosome arrays (Bustamante et al., 2021). This type of assay is able to reveal heterogeneous and long-range interactions between chromatin and chromatin-binding complexes, and quantify the strength of their interactions (Leicher et al., 2020). We reconstituted 12-mer nucleosome arrays harboring either canonical H2A or H2A.Z histone octamers. A single array is tethered between a pair of beads through two DNA handles (Figure 5A). Each bead is controlled by an optical trap, which allows us to pull on the tethered array by gradually separating the two beads. The resultant force-distance (f-d) curve displays transitions that correspond to the force-induced disruption of molecular contacts. Without SUV420H1, the f-d curve for both canonical H2A and H2A.Z arrays shows the signature sawtooth pattern in the high force regime (>15pN) (Brower-Toland et al., 2002; Leicher et al., 2020) representing the abrupt unraveling of individual nucleosome inner wraps, as well as a plateau in the low force regime (<10pN) corresponding to the unwrapping of the outer turn of nucleosomal DNA along with disruption of inter-nucleosomal stacking interactions (Figures 5B and S13A) (Meng et al., 2015). The presence of SUV420H1 drastically changes the f-d curves for both types of arrays. We observed that transitions start to occur at lower forces, disrupting the low-force plateau typically seen on bare nucleosome arrays, indicating that the outer wrap of the nucleosomal DNA is engaged with SUV420H1 (Figures 5B and S13A). We also observed larger transitions in the f-d curves for SUV420H1-bound arrays, indicating that SUV420H1 may mediate inter-nucleosome interactions. Overall, the presence of SUV420H1 yields a broader distribution of transition forces (Figures 5C and S13B), consistent with the view that this complex physically alters the structural organization of chromatin. The ΔC-terminus mutant, on the other hand, did not noticeably alter the f-d curves of chromatin arrays, resembling that of the f-d curves of chromatin alone without SUV420H1 (Figure S13C). Generally, wildtype SUV420H1 induces transitions at lower forces (<10pN) which is not observed with the ΔC-terminus mutant or chromatin alone (Figure S13D). SUV420H1 does not interact with bare DNA in the same manner as chromatin, illustrated by optical trap control experiments with wildtype and ΔC-terminus SUV420H1 against bare DNA (Figure S13E). In line with this data, SUV420H1 contributes to heterochromatin formation *in vivo* (Bromberg et al., 2017; Jorgensen et al., 2013; Schotta et al., 2004). H4-K20 trimethylation is strongly enriched at condensed pericentric heterochromatin and at inactive X chromosome (Jorgensen et al., 2013). We wondered whether SUV420H1 has chromatin condensation activity similar to HP1 since both of them cooperate at heterochromatin loci. To test this idea, we employed an *in vitro* assay using fluorescently labeled H3K9me3 nucleosome arrays to which we added increasing amounts of SUV420H1. We observed that the chromatin arrays start to coalesce to form irregular phases at a SUV420H1 concentration of 0.625μM (Figure 5D). This *in vitro* condensation and documented heterochromatic localization of Suv420 proteins prompted the exploration of chromatin condensation by SUV420H1 in the presence of HP1α. In the *in vitro* assay with the labeled H3K9me3 nucleosome array, we introduced HP1α at a fixed concentration to which we added increasing amounts of SUV420H1. Without SUV420H1 present, HP1α promoted the formation of spherical condensates consistent with earlier reports (Larson et al., 2017; Sanulli et al., 2019; Strom et al., 2017). With an increasing amount of SUV420H1, droplets were driven towards a more irregular shape suggestive of changes in material properties (Figure 5E). These results suggest that SUV420H1 alone and in cooperation with HP1 may drive chromatin condensation in the context of heterochromatin.

**Figure 5.**
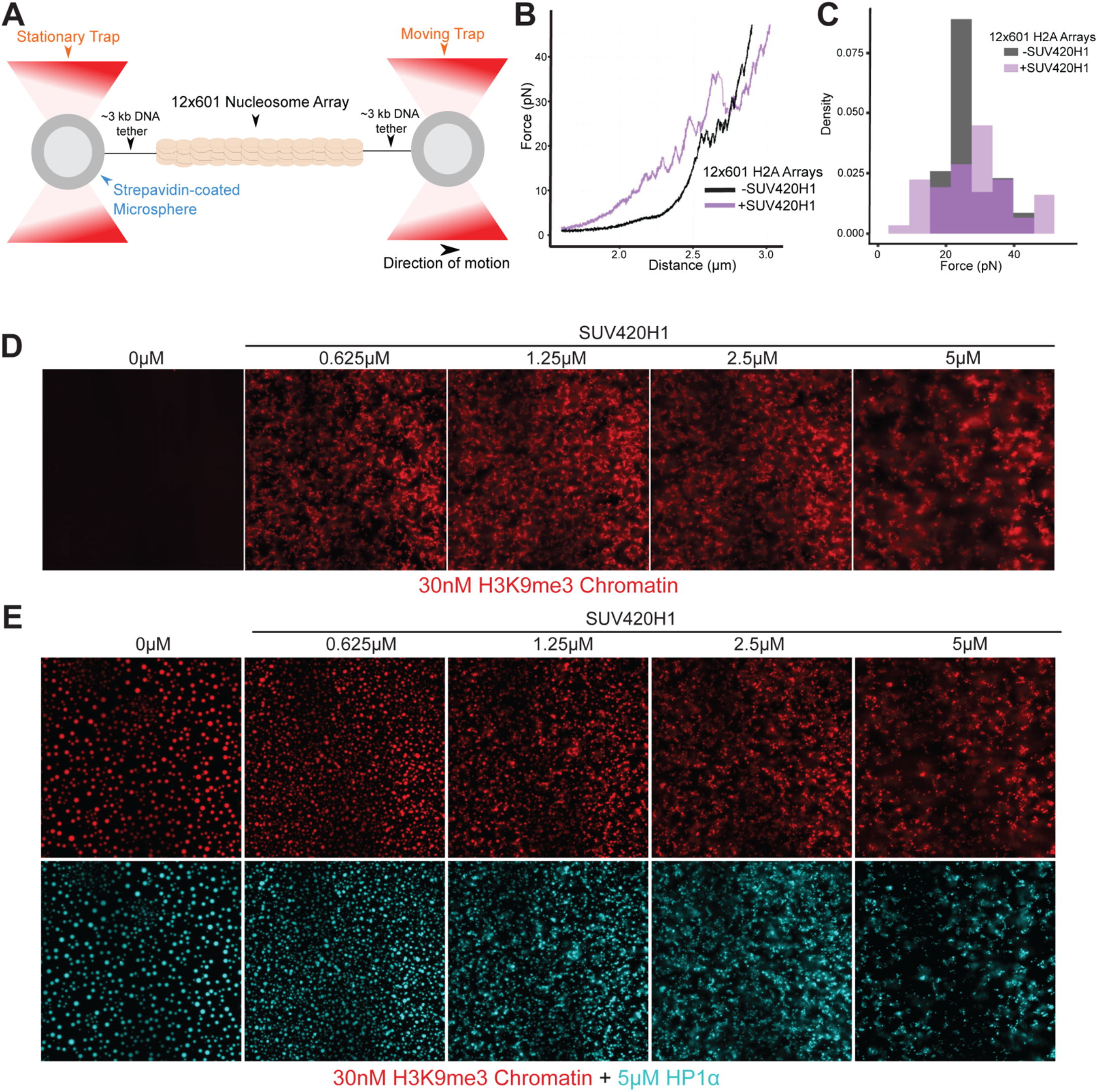
Biophysical and chromatin condensation activity of SUV420H1. A. Schematic representation of the optical tweezers setup where a doubly biotinylated 12-mer nucleosome array with long DNA handles is tethered to a pair of streptavidin-coated beads. B. A representative force-extension curve for an unbound histone H2A 12-mer array (black) and one for a SUV420H1- bound array (purple). C. Distribution of transition forces observed in the force-extension curves in (B). D. Micrograph panel of H3K9me3 chromatin array mixed with serial dilutions of SUV420H1 showing chromatin condensation. E. Fluorescence microscopy panel of H3K9me3 chromatin array (red) with HP1α (cyan) and serial dilutions of SUV420H1

### SUV420H1-nucleosome interactions in cells

To study how interactions of SUV420H1 with nucleosomes affect enzymatic activity in a physiological context, we introduced wildtype and various mutants of SUV420H1 into Suv4-20h double-null mouse embryonic fibroblasts (MEFs).

Using western blotting, we assessed the effects of the different mutations on H4K20 methylation (Figure 6A). As expected, there is a global loss of H4K20me2/3 in the Suv4-20h double-null MEFs cells and the catalytically dead mutant (S251A), however, there was no apparent difference between wildtype and ΔC-terminus constructs. Consistent with our methyltransferase assays, we found that the DNA-interacting loop residue R286A reduces enzymatic activity. We also examined H4K20 methylation in MEFs using quantitative mass spectrometry (Figure 6B). In agreement with our western blot and enzymatic analyses, methylation levels of wildtype and ΔC-terminus SUV420H1 constructs were comparable in contrast to the R286A mutant, emphasizing that the C-terminal residues have minimal influence on catalytic activity. Suv420 null MEFs cells are hypersensitive to DNA damaging agents (Schotta et al., 2008). To test whether this attribute was a consequence of the catalytic or non-catalytic role of SUV420H1, we measured the sensitivity of cells expressing wildtype SUV420H1 or its mutants to DNA damage. To do this, we quantified cell proliferation (Crystal violet staining) after inducing DNA damage using etoposide (inhibitor of topoisomerase II). Indeed, we observed increased sensitivity of SUV420 KO to etoposide compared to Suv420H1 rescue cells. Compared to the wildtype, there was increased sensitivity to etoposide in the DNA interacting loop mutant (R286A), catalytically inactive mutant (S251A), and ΔC- terminus mutants (Figure S14A). To understand SUV420H1 catalytic and non-catalytic roles in cellular sensitivity to DNA damage, we examined the formation of P53 Binding Protein 1 (53BP1) foci in MEFs. Cells were exposed to etoposide, and the nuclear localization of 53BP1 post-exposure was examined. In contrast to the knockout and SUV420H1 mutants (R286A and ΔC-terminus), cells with wildtype SUV420H1 exhibited a more robust response as determined by the percent of nuclear 53BP1 foci in the first couple hours of recovery (Figures 6C and S14B). The decrease in DNA damage response and cell survival to etoposide treatment suggests that the catalytic and the non-catalytic roles are important to SUV420H1 function in DNA damage response.

**Figure 6.**
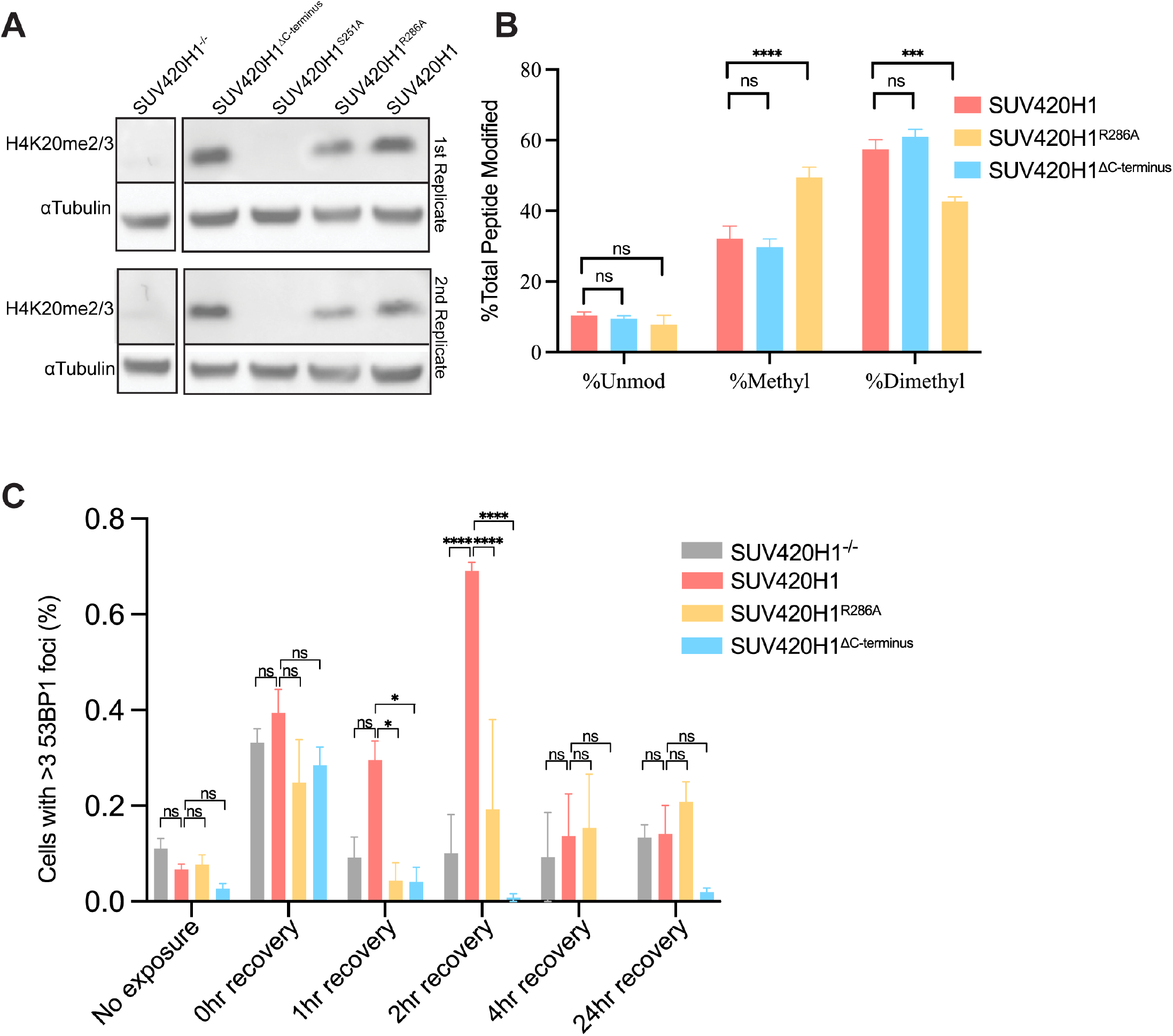
Cellular activity of SUV420H1. A. Representative western blot showing the level of H4K20me2/3 in Suv420H1 MEFs, loading control αTubulin. B. Relative abundance of histone H4 lysine 20 methylation in wildtype, R286A and ΔC-terminus Suv420H1 containing MEFs. Bars graphs represent the means ± SEMs of four independent biological replicates. A 2way ANOVA statistical analysis was performed (*p < 0.05, ***p < 0.001, ****p < 0.0001) C. Percentage of cells with >3 53BP1 foci per nucleus in SUV420H1 knockout, wildtype, R286A and ΔC-terminus MEFs cells. Bars graphs represent the means ± SEMs of three independent biological replicates. A 2way ANOVA statistical analysis was performed (*p < 0.05, ****p < 0.0001)

## Discussion

We provide the structural and biochemical characterization of SUV420H1 interaction with H2A.Z and H2A-containing nucleosomes. Our studies uncover the interfaces between SUV420H1 and nucleosomes and shed light on the mechanisms of several cancer-associated mutations. Our work provides a framework to understand how SUV420H1 catalytic activity is enhanced on H2A.Z-containing nucleosomes due to an extended nucleosome acidic patch. The cryo-EM maps of the complexes reveal a non-catalytic activity of SUV420H1 that causes a detachment of terminal DNA from the histone octamer. We show that the detachment is at least in part due to the C-terminal portion of SUV420H1, which in the structure can be found close to the αN helix of histone H3. This observation points to a possible role of SUV420H1 in eukaryotic genomes, where SUV420H1 binding to chromatin could influence DNA accessibility. Increased DNA accessibility, particularly in DNA repair and replication, is required for recruiting DNA-associating factors (Groth et al., 2007; Hauer and Gasser, 2017; Zhou et al., 2019). It could also potentially enable access to sites on nucleosomes that, under physiological conditions, are inaccessible due to DNA–histone interactions, as observed in canonical nucleosomes. One example here would be HP1, which interacts with the H3 αN-helix region and has recently been shown to probe the buried surface of the nucleosome core (Dawson et al., 2009; Sanulli et al., 2019). One of the notable surfaces exposed upon DNA detachment, as seen in our SUV420H1 bound nucleosome models, is the α1-helix of H2B, which contains a canonical HP1 interacting PxVxL sequence motif. We hypothesize that the activity of SUV420H1 in exposing buried histone residues would facilitate HP1 binding to chromatin and chromatin condensation. This would help explain the studies on HP1 interactions with nucleosome surfaces engaged in histone: DNA interactions within nucleosomes. (Dialynas et al., 2006; Jang et al., 2014; Lavigne et al., 2009; Sanulli et al., 2019).

We demonstrate that SUV420H1 alone could contribute to chromatin condensation. Our observations regarding chromatin condensation further rationalize the roles of SUV420 proteins at heterochromatin (Jorgensen et al., 2013; Schotta et al., 2004). Our *in vitro* work also shows that SUV420H1 changes the conformation of chromatin, consistent with the functional importance of the αN helix of histone H3 in regulating chromatin dynamics. Our structures show that increased interaction between the C-terminus of SUV420H1 and the αN helix of histone H3 correlates with DNA detachment, influencing nucleosome breathing. Proteins that work in a wide array of chromatin-dependent functions, such as transcription, rely on transient access to DNA and are made possible by nucleosome breathing. Recently, it was reported that nucleosome breathing also enhances nucleosome multivalency dictating chromatin conformation (Farr et al., 2021). Another explanation linking nucleosomal DNA detachment and chromatin condensation activities would be that allowing more flexibility of the DNA ends allows chromatin to adopt more compact structures, as previously suggested for the role of H2A.Z (Lewis et al., 2021). The influence breathing has on nucleosome arrays’ self-association property plays a fundamental role in the nature of the genome and its proposed ‘liquid-like’ state, as well as the formation of distinct genomic compaction and organization (Gibson et al., 2019).

SUV420H1 role in the genome is multifaceted, from its catalytic function being histone H4K20 methylation to its non-catalytic function in DNA detachment, with a consequential role in chromatin dynamics by increasing nucleosomal breathing. The uncoupling of catalytic and non-catalytic roles of histone-modifying proteins/complexes has taken center stage in recent years. The catalytically independent functions discovered in PRC1, PRC2, COMPASS/Set1A, and yeast Dot1, to name a few, have emerged as essential components of their function (Grau et al., 2021; Illingworth et al., 2015; Lee et al., 2018; Morgan and Shilatifard, 2020; Sze et al., 2017). Our results provide a structural and biochemical basis for both the catalytic and non-catalytic roles of SUV420H1 in genome regulation and open new avenues for studying these enzymes and their function.

## Material and Methods

### SUV420H1 expression and purification

Human SUV420H1, its mutants, and ΔC-terminus (residues 362-393 deleted) protein construct were cloned into a pGEX- 4T2 vector with an N-terminal GST-tag with a thrombin cleavage site. The expression and purification of SUV420H1 and mutants were performed using the same protocol. SUV420H1 plasmid was transformed into *Escherichia coli* BL2-codon plus (DE3)-RIL competent cells (Agilent technologies) and grown in 2xYT-Kan media. At an OD_600_ between 0.4–0.6, cell cultures were induced with 0.5 mM IPTG at 18 °C for an overnight expression. The cells were centrifuged at 4,000 rpm for 30 mins at 4 °C (Sorvall LYNX6000) and lysed (AvestinEmulsiflexC3) with lysis buffer (20 mM Tris, pH 7.9, 300 mM NaCl, 5% Glycerol, 1 mM ß-mercaptoethanol (BME), 1x protease inhibitor cocktail (Roche)). The slurry was then centrifuged at 19,000 rpm for 30 mins, and the soluble fraction was collected. SUV420H1 proteins were isolated by incubating the soluble fraction for 2 hrs with GST beads (Macherey-Nagel). The beads were washed with two column volumes of wash buffer (20 mM Tris, pH 7.9, 300 mM NaCl, 1 mM BME) and one column volume of wash buffer containing high salt (2 M NaCl). Protein was released from the GST beads by thrombin cleavage and purified by MonoQ (GE Healthcare) ion-exchange chromatography. Following ion exchange, the fractions containing SUV420H1 were purified on HiLoad Superdex 200 10/300 (GE Healthcare) size-exclusion chromatography column (S200 Buffer: 300 mM NaCl, 50 mM Tris pH 7.9, 5% Glycerol, 2 mM BME). Peak fractions were pooled, concentrated (∼4 mg/ml) and flash-frozen in liquid nitrogen and stored at -80 °C.

### Preparation and reconstitution of nucleosomes

Plasmids containing wildtype histones, and polycistronic histone plasmid were generous gifts from Drs. Karolin Luger and Peter Jones. Histone H2A chimeric mutant mimicking H2A.Z acidic patch (N94D/K95S) was generated using the Q5 mutagenesis kit (New England Biolabs). Monomethylated histone H4 lysine 20 (H4K20me1) was generated by alkylation of H4K20C following an established methyl lysine analog protocol(Simon, 2010). Expression and purification of wild-type individual histones and polycistronic or mutant polycistronic histones were conducted following the standard protocols as described(Armache et al., 2011; Xu et al., 2020). Briefly, individual histones were expressed in Rosetta (DE3) cells (Novagen), extracted from inclusion bodies, and purified by size exclusion and anion chromatography using previously published protocols(Dyer et al., 2004). Purified histones were freeze-dried using a Sentry lyophilizer (VirTis). For wildtype polycistronic and mutant polycistronic histone, a plasmid was transformed into *E.Coli* BL21(DE3) pLysS cells for large-scale expression. Cell pellets were lysed and supernatant purified through Ni-NTA agarose beads (QIAGEN) (Lysis Buffer: 20 mM Tris-HCl, pH 8.0, 2.0 M NaCl, 25 mM imidazole, 10% glycerol, 1 mM phenylmethanesulphonyl fluoride (PMSF) and 0.5 mM tris(2-carboxyethyl) phosphine (TCEP)/Elution Buffer: 20 mM Tris-HCl, pH 8.0, 2.0 M NaCl, 250 mM). His6-Sumo tags were cleaved by incubation with purified ULP1 Sumo protease at a 1:1,000 ratio at 4 °C overnight. Octamer was subsequently followed by further purification by Superdex 200 16/600 column (GE Healthcare) size-exclusion chromatography column (S200 Buffer: 10 mM Tris-HCl, pH 8.0, 2.0 M NaCl, 1mM EDTA, and 0.5 mM TCEP). Peak fractions were pooled, and concentrated.

To generate nucleosomal DNA, a plasmid comprising eight copies of the Widom 601 nucleosome positioning sequence(Lowary and Widom, 1998), each flanked by the EcoRV restriction enzyme cutting site(Armache et al., 2011), was transformed into DH5a (ThermoFisher) competent cells and grown in 2xYT-Amp media overnight. The 601 DNA fragment was excised using EcoRV and purified using previously published protocols(Dyer et al., 2004).

Nucleosome reconstitutions from individual histones were done as described(Armache et al., 2011; Dyer et al., 2004; Xu et al., 2020). Briefly, equimolar amounts of H3, H4, H2B, and H2A or H2A.Z were mixed and dialyzed into refolding buffer (2 M NaCl, 10mM Tris-HCl pH 7.5, 1 mM EDTA, 5 mM DTT). The octamer in refolding buffer was purified by size exclusion chromatography on a Superdex 200 16/600 column (GE healthcare). Nucleosomes were assembled by incubating purified Widom 601 DNA and histone octamers assembled from individual histone or polycistronic histones and dialyzed overnight with gradient salt dialysis using a peristaltic pump (Gilson Rapid Pump). Reconstitution was completed by dialyzing into TCS buffer (20 mM Tris-HCl pH 7.5, 1 mM EDTA, 1 mM DTT), and reconstitution efficiency was analyzed by 4.5% PAGE gel and quantified based on the DNA content.

### Endpoint methylation assay

SUV420H1 methyltransferase activity was determined by monitoring the production of S-adenosyl homocysteine (SAH) in the presence of S-adenosyl methionine (SAM) (Sigma) and nucleosomes. The wild-type or mutant SUV420H1 at concentrations of 50 or 100 nM was incubated with 1 µM H2A and H2A.Z nucleosomes with a methyltransferase buffer (20 mM Tris pH 8.0, 50 mM NaCl, 1mM DTT, 0.1 mg/mL BSA, 40 µM SAM). 20 µL methyltransferase reactions were performed at 30°C for 15 mins and stopped with 5 µL of 0.5% trifluoroacetic acid (TFA). The SAH production was measured using MTase-Glo methyltransferase kit (Promega) using 8 µL of the stopped reaction. 2 μL MTase-Glo Reagent was added to 8 uL of the stopped reaction to convert SAH to ADP, and 10 μL MTase-Glo Detection solution was added to transform ADP to ATP. The luminescence of the product was measured using an EnSpire 2300 Multilabel plate reader (Perkin Elmer). All assays were performed with three replicates.

### Kinetic assays

To measure the *k*_cat_ and *K*_M_ values for SUV420H1 activity on nucleosomes, nucleosomes containing either H2A or H2A.Z were 2-fold serially diluted from 4000 nM – 4 nM. Additionally, a stock solution of SUV420H1 at 20 nM containing 40 μM SAM (Sigma) was prepared with a reaction buffer (20 mM Tris pH 8.0, 50 mM NaCl, 1mM DTT, 0.1 mg/ml BSA). To initiate the reaction, 10 μL of SUV420H1 and 10 μL of nucleosomes were incubated at 30°C for 15 mins. The reaction was quenched with 5 μL 0.5% TFA. The SAH production was determined with MTase-Glo methyltransferase kit (Promega) using 8 uL of the stopped reaction. 2 μL MTase-Glo Reagent was added to 8 µL of the stopped reaction, and 10 μl MTase-Glo Detection solution was added. The luminescence of the product was measured using an EnSpire 2300 Multilabel plate reader (Perkin Elmer). A linear regression fit of the SAH calibration curve was determined and served to interpolate SAH produced from the reactions. Kinetic parameters for methyltransferase activity were determined using enzymatic kinetics analysis in Prism 8 (GraphPad). All assays were performed with three replicates.

### Western blots for in vitro methyltransferase assay

Methylation reactions were performed by incubating SUV420H1 at concentrations of 50 and 100 nM with 1 µM H2A and H2A.Z containing H4K_C_20me1 nucleosomes in a methyltransferase buffer (20 mM Tris pH 8.0, 50 mM NaCl, 1 mM DTT, 0.1 mg/mL BSA, 40 μM SAM). With a reaction volume totaling 20 µL, methylation reaction was performed at 30°C for 15 mins and quenched with 5.0 µL 5X SDS buffer and resolved by electrophoresis using a 15% SDS-PAGE gel, and transferred into 0.2 mm polyvinylidene difluoride PDVF membrane (Bio-Rad). H4K20 methylation was determined by incubating the membranes with anti-H4K20me1 (Abcam ab9051, 1:1000), anti-H4K20me2 (Abcam ab9052, 1:1000) and anti- H4K20me3 (Abcam ab17567, 1:1000) for 12 h at 4°C. The western blots were developed with an ECL reagent (Thermo Fisher) and imaged in the ChemiDoc system (Bio-Rad). The loading control was determined with FastBlue staining.

### Sample preparation using gradient fixation (GraFix)

All cryo-EM samples described in this study were prepared using the GraFix protocol(Stark, 2010). SUV420H1- H2A/H2A.Z nucleosomes, SUV420H1 C-terminal peptide-nucleosome complex and nucleosomes alone were dialyzed into Buffer A (50 mM HEPES pH 7.9, 100 mM NaCl, 5% Glycerol, 1 mM DTT) with molar ratio of 1:5 (Nucleosome:SUV420H1) or 1:100 (Nucleosome: SUV420H1 C-terminal peptide) in the presence of 40 µM of SAM (Sigma). Samples were incubated on ice for 20 mins. Gradient was made with buffer B (50 mM HEPES pH 7.9, 100 mM NaCl, 10% Glycerol, 1 mM DTT) and buffer C (50 mM HEPES pH 7.9, 100 mM NaCl, 5% Glycerol, 1 mM DTT, 0.1% glutaraldehyde) using a gradient maker (Biocomp gradient master). The sample was run on the gradient for 16 hrs at 4 °C using an Optima XE-90 ultracentrifuge (SW40Ti rotor, Beckman-Coulter) at 30,000 rpm. The resultant gradient was then fractionated and analyzed via gel electrophoresis. Selected fractions (Figure S2) were dialyzed into buffer containing 20 mM HEPES pH 7.9, 100 mM NaCl, 2 mM DTT, and concentrated for grid freezing.

### Cryo-EM sample preparation, data acquisition, and processing

Cryo-EM grids of the SUV420H1 in complex with H2A or H2A.Z nucleosome, SUV420H1 C-terminal peptide in complex with H2A.Z nucleosome, and H2A/H2A.Z nucleosome alone were prepared following established protocol(Li et al., 2013). 3.0μL of the samples with concentrations in the range of 0.1 to 0.6 mg/mL were applied to Quantifoil gold grids (300 mesh, 1.2 μm hole size) glow-discharged for 25s. The grid was blotted for 3s with a blot force of 3 using Vitrobot Mark IV (FEI Company) at 4 °C and 100% humidity and plunge frozen in liquid ethane. Data for SUV420H1-H2A/H2A.Z complexes were collected on an FEI Titan Krios 300 kV equipped with a Gatan K3 Summit camera at a nominal magnification of 81,000× and an image pixel size of 1.08Å. Images were collected as a movie stack for 3.5s, fractionated into 40 subframes, and the accumulated dose of 50 e^−^/Å^2^. For the SUV420H1-H2A.Z complex, a total of 5445 images were collected at a defocus range of −1 µm to −2.4 µm, and for the SUV420H1-H2A complex, 8004 images were collected at a defocus range of −0.25 µm to −1.75 µm. The movies were motion-corrected and binned to 1.08 Å/pixel using UCSF MotionCor2 v1.4.7(Zheng et al., 2017). The Contrast Transfer Function (CTF) was calculated using cryoSPARC’s ‘Patch CTF Estimation (multi)’; Particles were picked with the ‘Blob Picker’ with a diameter between 100- 140Å, and were extracted from the micrographs in a 256 x 256 box, Fourier cropped to box 96 x 96 for the initial processing steps(Punjani et al., 2017). Particles were subjected to 2D classification into 50 classes, and the best particles were used to generate 3 models using the ‘Ab initio’ function. We selected the best map from the ‘Ab initio’ function reflecting nucleosome features and put it through 3D classification using all the particles extracted (5,506,853 for SUV420H1-H2A.Z complex and 4,761,686 SUV420H1-H2A complex). We then selected the best subset, refined it and re-extracted particles in a 256 x 256 box. These particles were further classified, refined and polished to obtain the final reported maps. SUV420H1-H2A.Z complex was resolved at 3.08 Å from 459,864 particles, and was subsequently subjected to further optimization using Bayesian Polishing, which improved the resolution to 2.63 Å; SUV420H1-H2A complex yielded a resolution of 3.37 Å from 366,390 particles.

H2A.Z/H4K20M nucleosome alone was collected on FEI Titan Krios 300kV equipped with a Gatan K3 collected at 105,000× magnification (calibrated pixel size of 0.4260Å). Movies were collected using Leginon (Suloway et al., 2005) with a total exposure of 3.00 seconds, for an accumulated dose of 54.76 e^-^/Å^2^ fractionated over 50 frames. A total of 894 images were collected at a defocus range of 1.3 – 1.5 μm. Movie stacks acquired in counting mode were corrected for global and local motions (in 5x7 patches) using UCSF MotionCor2 v1.2.1(Zheng et al., 2017). Image processing steps were carried out only in cryoSPARC using an approach applied to SUV420H1-H2A/H2A.Z complexes. This processing scheme resulted in a final reconstruction of the H2A.Z/H4K20M nucleosome at a resolution of 3.07Å from 206,872 particles.

Cryo-EM dataset for H2A nucleosome alone and SUV420H1 C-terminal peptide-H2A.Z nucleosome complex was collected on an FEI Talos Arctica 200 kV equipped with a Gatan K3 Summit camera. Images were collected at a magnification of 36,000× with a calibrated pixel size of 0.5480Å used for processing. Movies were collected using Leginon at a dose rate of 21.20 e^-^/Å^2^/s with a total exposure of 2.40 seconds, for an accumulated dose of 50.89 e^−^/Å^2^ fractionated over 48 frames. 175 and 846 images were collected for H2A nucleosome alone and SUV420H1 C-terminal peptide-H2A.Z nucleosome complex, respectively. Movie stacks acquired in Super-resolution mode were corrected for global and local motions (in 5x7 patches) using UCSF MotionCor2 v1.2.1. Image processing steps were carried out only in cryoSPARC using an approach applied to SUV420H1-H2A/H2A.Z complexes. The best-resolved maps presented overall resolutions for H2A nucleosome alone, and SUV420H1 C-terminal peptide-H2A.Z nucleosome complex was 3.86Å from 79,392 particles and 3.08Å from 494,524 particles respectively. The final resolutions reported for all reconstructions were determined using Fourier Shell Correlation (FSC) at 0.143 cutoffs(Rosenthal and Henderson, 2003) following gold-standard refinement. For comparison, SUV420H1-H2A.Z and SUV420H1-H2A cryoEM maps were sharpened using the DeepEMhancer in cryoSPARC with tightTarget model setting (Sanchez-Garcia et al., 2021). The processing details and summaries are shown in Figures S3, S4, and S11

### Model building

The best-resolved, primary cryo-EM map of the SUV420H1-H2A.Z complex at 2.63 Å Coulomb potential density allowed unambiguous fitting of SUV420H1, histones proteins, and DNA. The pre-polished final map at 3.08 Å from cryoSPARC was used to reconstruct the C-terminal region of SUV420H1. Available X-ray structures were rigid-body-fit into cryo-EM reconstructions for the catalytic domain of SUV420H1 (PDB:3S8P), H2A.Z nucleosome (PDB: 1F66), and H2A nucleosome (PDB: 1AOI)(Luger et al., 1997; Suto et al., 2000; Wu et al., 2013). These PDBs were first manually fit into the density and then locally optimized using UCSF Chimera’s "Fit in map" function(Pettersen et al., 2004). We then used Coot(Emsley and Cowtan, 2004) for local adjustments of secondary elements and sidechains into densities. The C-terminal of SUV420H1 was built de novo manually using Coot. The complete model was refined using PHENIX, using first only rigid body setting and then secondary structure, ADPs, rotamer, and Ramachandran restraints in 100 iterations(Adams et al., 2010). Ramachandran outliers and regions that needed manual fixing were attended to in Coot. Chimera and PyMOL were used to prepare figures of the model and cryo-EM densities(Emsley and Cowtan, 2004; Pettersen et al., 2004; Schrodinger, 2015). Final model statistics are reported in Extended Data Table 2.

### SUV420H1-H2A.Z nucleosome complex crosslinking for mass spectrometry

SUV420H1-H2A.Z nucleosome complex was assembled in a ratio of 3:1 and was purified on HiLoad Superdex 200 10/300 (GE Healthcare) size-exclusion liquid chromatography column (Buffer: 100 mM NaCl, 50 mM HEPES pH 7.5, 1mM DTT) Peak fractions representing the complex were pooled and concentrated to 0.5 mg/mL. Amine-specific crosslinking of histones within the complex was done with bis(sulfosuccinimidyl)suberate (BS3, ThermoFisher Scientific). 20-μl reactions containing complex in (20 mM HEPES pH 7.5, 100 mM NaCl, and 1 mM DTT) and 0.5 to 2 mM BS3 were incubated on ice for 2 hours before being quenched with Tris-HCl pH 7.5 added to 50mM from 0.5 M stock. After 10min incubation with Tris, proteins were precipitated with acetone: 4 volumes of acetone (cooled at -20 °C) were added, followed by 1-hour incubation at -20 °C. Protein precipitates were collected by 10-min centrifugation at 16000×g at room temperature. Pellets were rinsed with 80% acetone, dried on-air, and dissolved in 9 μL of 50mM ammonium bicarbonate, 10mM DTT, and 8 M urea buffer. After 10mins of incubation at room temperature, samples were mixed with 1 μL 0.5 M iodoacetamide and incubated for 30mins in the dark. For digestion, 10μl samples were diluted with 100 μL 50 mM ammonium bicarbonate, 5mM DTT containing 10 ng/μL trypsin/Lys-C mixture (Promega), and incubated overnight at 25 °C. Digestion reactions were stopped by mixing with 10 μL 20% heptafluorobutyric acid and clarified by 10 mins centrifugation at 16000×g. Finally, peptides were desalted using C18 OMIX tips (Agilent) according to the manufacturer’s protocol, dried under vacuum, and dissolved in 15 μL 0.1% formic acid.

### Mass spectrometry

Peptides were analyzed in the Orbitrap Fusion Lumos mass spectrometer (Thermo Scientific) coupled to Dionex UltiMate 3000 (Thermo Scientific) liquid chromatography system. Peptides were resolved on a 50 cm long EASY-Spray PepMap RSLC C18 column using a 120mins linear gradient from 96% buffer A (0.1% formic acid in water) to 40% buffer B (0.1% formic acid in acetonitrile) followed by 98% buffer B over 5mins with a flow rate of 150nl/min. Each full MS scan (orbitrap analyzer, resolution 60,000) was followed by data-dependent MS/MS scans (orbitrap, resolution 15,000) for the 20 topmost abundant peptides after HCD fragmentation. The quadrupole isolation window was set to 2m/z, and precursors with charge states 4 to 6 were selected for fragmentation with collision energy set to 35. Monoisotopic precursor selection was enabled, and a dynamic exclusion window was set to 30 sec.

### Mass spectrometry data processing and analysis

Raw data was processed with pLink2(Chen et al., 2019) using a database of concatenated sequences of proteins constituting nucleosomes and common contaminants. For peptides identification, up to 3 missed trypsin cleavages were allowed, variable modifications were oxidation at methionine residue and carbamidomethyl at cysteine residue, cross- linker was set to BS3, FDR level was set to 1%, precursor and fragment tolerance were set to 10ppm and 50ppm, respectively. Other parameters were left unchanged. A list of identified crosslinked peptides was imported into the R environment for statistical computing and filtered to only those with reported Q-value below 0.01. Crosslinking sites confirmed by at least two peptide-spectrum matches were considered for further analysis.

### Nucleosome array reconstitution for optical tweezers assay

Twelve Widom 601 sites with 30bp linkers were inserted into a pTXB1 vector, and the resulting plasmid was subsequently digested with PciI to create DNA handles of approximately 3.2 and 3.4 kb on either side of the 12-mer 601 sequence. Biotins were then added to the ends of the digested plasmid via Klenow fill-in. Nucleosome arrays were reconstituted via the standard salt dialysis method(Dyer et al., 2004; Leicher R, 2022). Briefly, prepared DNA constructs, recombinant human histone octamers, and MMTV competitor DNA was mixed in high salt buffer (10 mM Tris-HCl pH 7.5, 1mM EDTA, 2 M NaCl, 5 mM beta-mercaptoethanol, 0.01% Triton X-100). The nucleosome array mixture was then slowly dialyzed via buffer exchange to the low salt buffer (10 mM Tris-HCl pH 7.5, 1 mM EDTA, 50 mM NaCl, 5 mM beta-mercaptoethanol, 0.01% Triton X-100) over the course of roughly 30-40 h. Exact concentrations of DNA and histones were determined via titrations for each round of reconstitutions but generally were within the range of 1:0.2:0.3 molar ratios of DNA construct: Histone Octamers: MMTV, respectively, with a final total 601 sequence concentration of 1 μM.

The quality of the reconstituted arrays was assessed via NspI digestion and through analysis of unwrapping peak numbers on the optical tweezers.

### Single-molecule optical tweezers assay

Force-distance curves for the 12-mer nucleosome arrays were collected using a LUMICKS C-Trap^®^ instrument(Leicher R, 2022; Meng et al., 2015). A single reconstituted arrays or bare DNA was tethered between a pair of 3.23μm streptavidin-coated beads (Spherotech) in a buffer containing 10 mM Tris-HCl pH 8.0, 50 mM NaCl, 0.5 mM MgCl_2_, 0.02% Tween-20, 0.01% BSA, and 1 mM DTT. The tethered arrays and bare DNA was then incubated with 10 nM SUV420H1 or ΔC-terminus mutant, and force-distance curves were obtained at a pulling speed of 50 nm/sec with trap stiffnesses around 0.6-0.7 pN/nm. The collected force data were down-sampled to 110Hz to match bead distance measurements for subsequent analysis. After a force-extension curve was recorded and accepted, the tether was pulled until it broke and a background trace was collected from the same pair of beads. The background force was then calculated and subtracted from each trace in the postprocessing steps. Peaks of force-induced transitions in the force-extension curve were identified using POTATO (Buck et al., 2021) using a z-score of 1.75, a median moving window size of 50, and a 5 pN minimum force threshold. Transitions that occurred above 50 pN were removed from analysis due to potential overlap with DNA overstretching peaks.

### In vitro phase-separation assay

SUV420H1-H3K_C_9me3 chromatin droplets were formed at 40nM nucleosome array concentration in 20 mM HEPES pH 7.5, 0.05 mM Tris pH 7.9, 0.05mM EDTA, 71 mM KCl, 14.3mM NaCl, 0.2% glycerol, 0.5 mM DTT, and 1 mM BME. The array DNA was composed of a fluorescent nucleotide Alexa Fluor 647-aha-dCTP (ThermoScientific) and was generated using Klenow Fragment 3’ – 5’ exo- (NEB). Nucleosome arrays were mixed with SUV420H1 or 5μM HP1α and incubated at room temperature for 20 mins before transferring to a glass bottom 384 well plate for imaging. 384 well glass bottom plates (Greiner Sensoplate 781892) were prepared for sample examination as follows: The wells were rinsed three times with 100 μL water, incubated with 100 μL of 2% Hellmanex for 30 mins-1 hr, rinsed with water three times again, incubated with 100 μL of 0.5 M NaOH for 30 mins, rinsed with water three times, and then 70 μL of 20 mg/ml mPEG-silane dissolved in 95% EtOH was added to coat the wells. The plate was covered with foil and left overnight at 4 °C. After the overnight, the wells were rinsed with 95% EtOH five times, followed by incubation with 100 mg/mL BSA for 2hrs. Then, the wells were rinsed three times with water and three times 2 with 1X phasing buffer afterwards (20 mM HEPES pH 7.2, 75 mM KCl, 0.05 mM EDTA, 0.5 mM DTT). After, the condensates are added to each well.

### Generation of mutant SUV420 cell lines

Human SUV420h1_i2 (2658bp) was amplified from human cDNA using PCR primers with XbaI and BamHI overhangs and cloned into XbaI/BamHI digested pCDH-EF1-MCS-Blasticidin (System Biosciences). Standard mutagenesis was used to generate the S251A, R286A, and ΔC-terminus mutations in SUV420h1_i2 with primers designed using NEBbaseChanger. To produce lentivirus, HEK293cells were transfected with the lentiviral vector (pCDH-EF1-blasticidin) carrying the SUV420H1 mutants and supporting plasmids (psPAX2 and pVSVG), followed by supernatant collection and filtering 72 h after transduction. SUV420 KO mouse fibroblast cells were cultured in Dulbecco’s modified Eagle’s medium (DMEM, Invitrogen) with 10% fetal bovine serum (FBS, Sigma). To generate transgenic expressing SUV420 mutants, SUV420 KO mouse fibroblast cells were transduced with a concentrated lentivirus. Transduced cells were grown under Blasticidin selection (6 μg/mL) 48 h after transduction and selected for one week.

Primers:

SUV420h1_i2_fw GCCTaCTCTAGAGTGGGTATGAAGTGGTTGG

SUV420h1_i2_Rev GGCCGCGGATCCTTACTTACTTGCATTGTTTTTTTC

R286A_Fw GTCAACTGGTgccGATACAGCATG

R286A_Rev ACAAACTTACAATTAGGTCTG

S251A_Fw AAACGACTTCgccGTCATGTACTCCAC

S251A_Rev TCTCCATGTCTAAGTAGC

C-terminus minus_fw TAAGGATCCGCGGCCGCG

C-terminus minus_Rev GTCACCTAACTTTTTAAGCCTATTTAAACGTTTATCTGTTTCTC

### Western blot for H4K20me2/3 using MEFs cells

Whole-cell lysates from 1 million cells were separated by SDS-PAGE (15 µg total protein), transferred to a polyvinylidene difluoride (PVDF) membrane, blocked with 5% nonfat milk in PBS containing 0.5% Tween-20 for 1 h at room temperature, probed with primary antibodies at a 1:1,000 dilution overnight at 4°C and detected with horseradish peroxidase-conjugated anti-rabbit or anti-mouse secondary antibodies at a 1:1,000 dilution. Primary antibodies: Anti-Histone H4 (dimethylK20,trimethyl K20) antibody [6F8-D9] (ab78517, abcam) and Anti-tubulin-alpha (9026,Sigma)

### Methylation of histone H4 in MEFs cells mass spectrometry

Histones were acid-extracted from MEFs cells as described (Shechter et al., 2007). Briefly, cell pellets (2 million cells per sample) were resuspended in NIB-250 buffer (15mM Tris, pH7.5, 60 mM KCL, 15 mM NaCl, 5 mM MgCl2,1 mM CaCl2, 250 mM sucrose, 1 mM DTT) and 0.3% NP-40, incubated for 5 min on ice, and centrifuged at 600 xg to collect the nuclei pellet. Pellets were washed with NIB two times to completely remove NP-40. Nuclei were resuspended in 0.4 N H2SO4 and rotate at 4 °C overnight. Samples were centrifuged at 3400 xg for 10 min and the supernatant was collected as the histone containing faction. Histones were precipitated with 25% trichloroacetic acid (TCA) overnight and washed with cold acetone. Samples were centrifuged at 3400 xg for 10 min and washed using acetone. Pellet was air dried overnight and suspended in 100 mM ammonium bicarbonate.

Histone samples were incubated with 100mM phosphate buffer and digested overnight with sequencing grade GluC (Roche) shaker at 37°C (500ng). 1/5 of digested sample was loaded onto a prepped EvoSep Tip for analysis. Tip was rinsed 3x with 50uL 0.1% Formic Acid and 100uL 0.1% Formic Acid was added to the tip before LC/MS/MS analysis. The run on Eclipse/EvoSep with 88min gradient/method that does HCD for all and additional ETD scans on precursors +3 and up between 400-800 m/z. LC separation online with MS using the autosampler of an Evosep. Peptides were gradient eluted from the column directly to the Orbitrap Fusion Eclipse (Thermo Scientific) using a 88 min gradient with Solvent A (Water, 0.1% formic acid) and solvent B (Acetonitrile, 0.1% acetic acid).

High resolution full MS spectra were acquired with a resolution of 120,000, an AGC target of 4e5, and scan range of 400 to 1500 m/z. All MS/MS spectra were collected using the following instrument parameters: resolution of 15,000, AGC target of 5e4, one microscan, 2 m/z isolation window, and HCD NCE of 30. An additional ETD MS/MS scan was taken precursors with charge states 3-15 between 400-800 m/z with the following parameters: resolution of 30,000, AGC target of 1e5, one microscan, 2 m/z isolation window, and ETD fragmentation with calibrated charge dependent parameters enabled. MS/MS spectra were searched using Byonic against Histone H4 and a Uniprot Mouse database. MS/MS spectra were also searched using Byos against just Histone H4 for quantitation and validation of peptides containing K20.

### Cell Proliferation Assay (Crystal violet staining)

Cells were collected, counted, and 1000 cells were seeded to a 12-well plate in quadruplicates in normal media (DMEM +10% FBS). The following day the cells were exposed to 0 µM (ctrl) or 1 µM etoposide for 3 h and then kept in culture for six days. The media was removed, and the cells were washed with PBS and stained with 0.5% Crystal Violet solution for 30 min at room temperature on a bench rocker, washed 4x with water, and dried overnight. 500 µL of methanol was added to each well and incubated for 20 min at room temperature on a bench rocker, and 200 µL was used to measure the absorbance at 570 nm in a 96 well plate.

### Immunofluorescence

53BP1 immunofluorescence cellular assay was performed as previously described (Taglialatela et al., 2021). SUV420 KO, SUV420H1, SUV420H1^R286A^ and SUV420h1^ΔC-terminus^ mouse fibroblast cells were collected, counted, and 6000 cells were seeded on black 96-well bottom-glass plates. After 24h the cells were exposed to 1 µM etoposide for 30 min and given 24h, 4h, 2h, 1h or 0h to recover after etoposide removal before the cells were simultaneously fixed and permeabilized (4% paraformaldehyde, 0.5% Triton X-100) for 10 min at room temperature. Cells were incubated in blocking solution (3% BSA in TBS-Tween 0.1%) for 1h with primary antibody (anti-53BP1, Rabbit polyclonal, Bethyl #A300-272A) diluted in blocking solution overnight at 4°C. After three washes with TBS-T the cells were incubated for 1 hr at room temperature with the secondary antibody Alexa Fluor 488-labeled anti-rabbit at 1:1,000 dilution (A-11008, Thermo Fisher Scientific). After three washes in TBS-T, cells were incubated with DAPI for 5 min at room temperature to stain nuclei. Image acquisitions were made using the ImageXpress Nano Automated Imaging System microscope (Molecular Devices).The imaging software MetaXpress 6 was used to automate image analysis. The experiment was done with three biological replicates for each clone.

## Acknowledgments

We thank Dr. William Rice, Dr. Alice Paquette, and Dr. Bing Wang for helping with data collection at NYU Langone Health cryo-EM Shared Resource. We thank the HPC Core at NYU Langone Health for computer access and support. This research was partly supported by the National Cancer Institute’s National Cryo-EM Facility at the Frederick National Laboratory for Cancer Research under contract HSSN261200800001E. We thank Peter Hare for his comments and critical review of this manuscript. We also thank Dr. Beatrix Ueberheide and Dr. Maria Antonelli for helping with mass spectrometric experiments at NYU Langone’s Proteomics Laboratory. The Proteomics Laboratory is supported by the NIH Shared Instrumentation Grant 1S10OD010582-01A1 for the purchase of an Orbitrap Fusion™ Lumos™ Tribrid™ mass spectrometer. We thank Dr. Gregory Bowman, Dr. Robert Kingston, Dr. Steven Henikoff, and Dr. Gunnar Schotta for fruitful discussions. We thank the Armache laboratory for critical comments and discussion. The work in the Armache laboratory is supported by the NIH grants R01GM115882, R01CA266978, and Mark Foundation for Cancer Research. S.A.-A. is supported by the Molecular Biophysics T32 grant (5T32GM088118). G.N., L.H., and T.L. are supported by the NIH R35 GM127020 and NSF-1921794 grants. S.L. is supported by the Robertson Foundation, the Pershing Square Sohn Cancer Research Alliance, the Starr Cancer Consortium, and an N.I.H. Director’s New Innovator Award (DP2HG010510). R.M.S. is supported by a National Science Foundation Graduate Research Fellowship. E.N. and N.V. are supported by the N.I.H. grant R01 GM127267, Blavatnik Family Foundation, and the Howard Hughes Medical Institute. The work in C.L.’s laboratory is supported by the NIH grant R01CA266978.

## Author contributions

S.A.A and K.-J.A. conceptualized and designed the study. S.A.A., P.D.I., R.L., and M.W. conducted structural and biochemical experiments. R.M.S and S.L. performed optical tweezers experiments. L.H., T.L., and G.N. performed the chromatin condensation assay. J.-P. A advised on cryo-EM analyses. M.D.S generated MLA histones, and N.V. and A. N. performed XLMS experiments. All authors contributed to data analysis, interpretation, and writing of the manuscript.

## Lead contact

All inquiries for further information and requests for resources and reagents should be directed to and will be fulfilled by the lead contact

---

**Figure S1.**
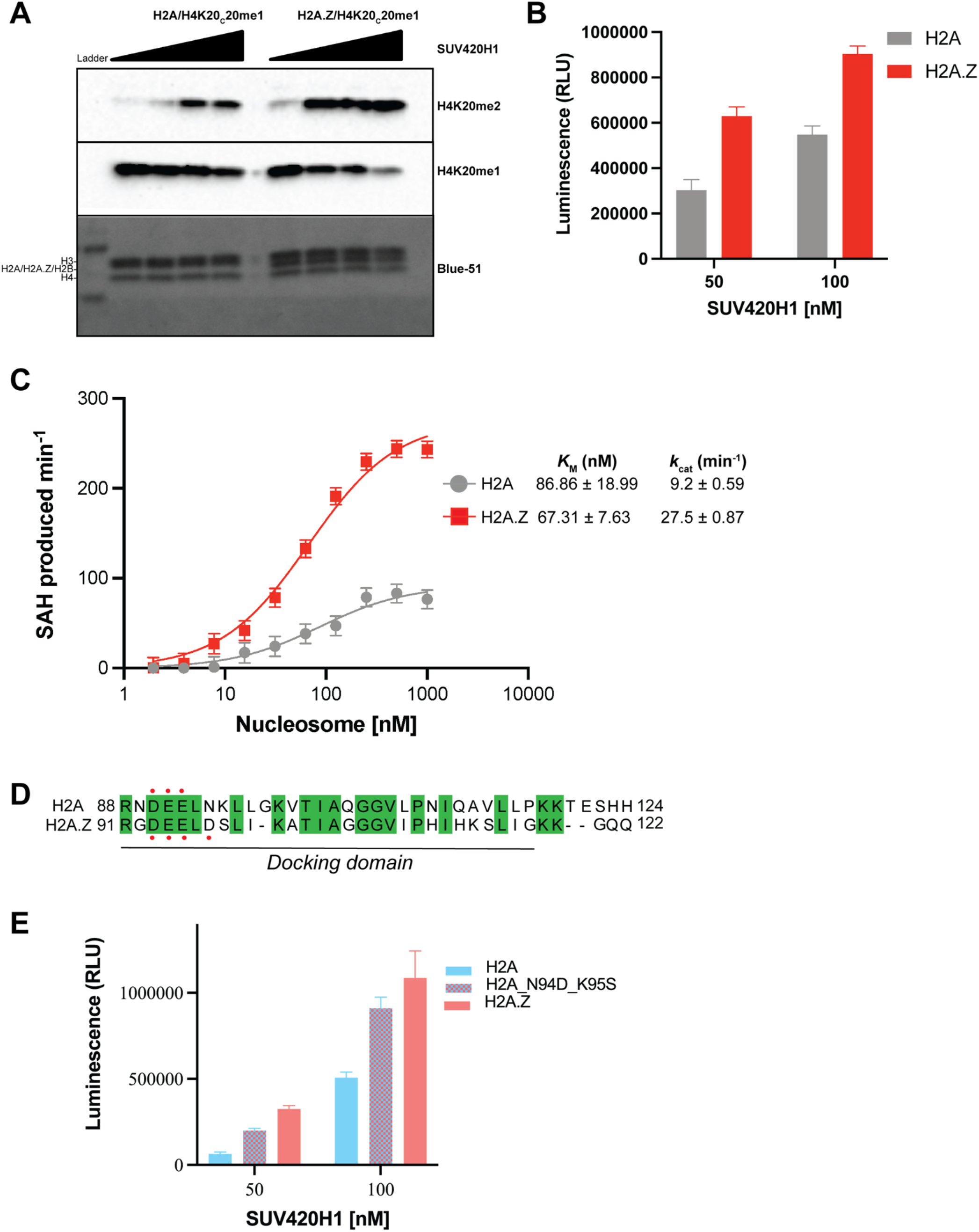
A. Representative western blot showing histone methylation assay measuring activity of SUV420H1 on H2A and H2A.Z nucleosomes. Methylation assays were performed with an increasing amount of SUV420H1 in the presence of H4K_C_20me1 nucleosome substrates. Reaction products were determined with antibodies against the different methylation states of H4K20. B. Catalytic activity of SUV420H1 on H2A and H2A.Z nucleosomes measured using an endpoint methyltransferase assay. The bar graph and error bars for the methyltransferase assay represents the mean ± SD from three independent experiments. C. Michaelis–Menten titrations of nucleosomes containing H2A and H2A.Z with SUV420H1. The *K*_M_ and *k*_cat_ values of the fitted data are reported in the graph. Each data point represents the mean ± SD from experiments repeated at least three times. D. Sequence alignment of the C-terminal domain of histones H2A and H2A.Z. Conserved residues are highlighted in green, and red dots represent the nucleosome acidic patch residues in the canonical H2A and the variant H2A.Z. E. Catalytic activity of SUV420H1 on H2A and chimeric H2A acidic patch mutant (N94D/K95S) nucleosomes measured using endpoint methyltransferase assay. The bar graph and error bars for the methyltransferase assay represents the mean ± SD from three independent experiments.

**Figure S2.**
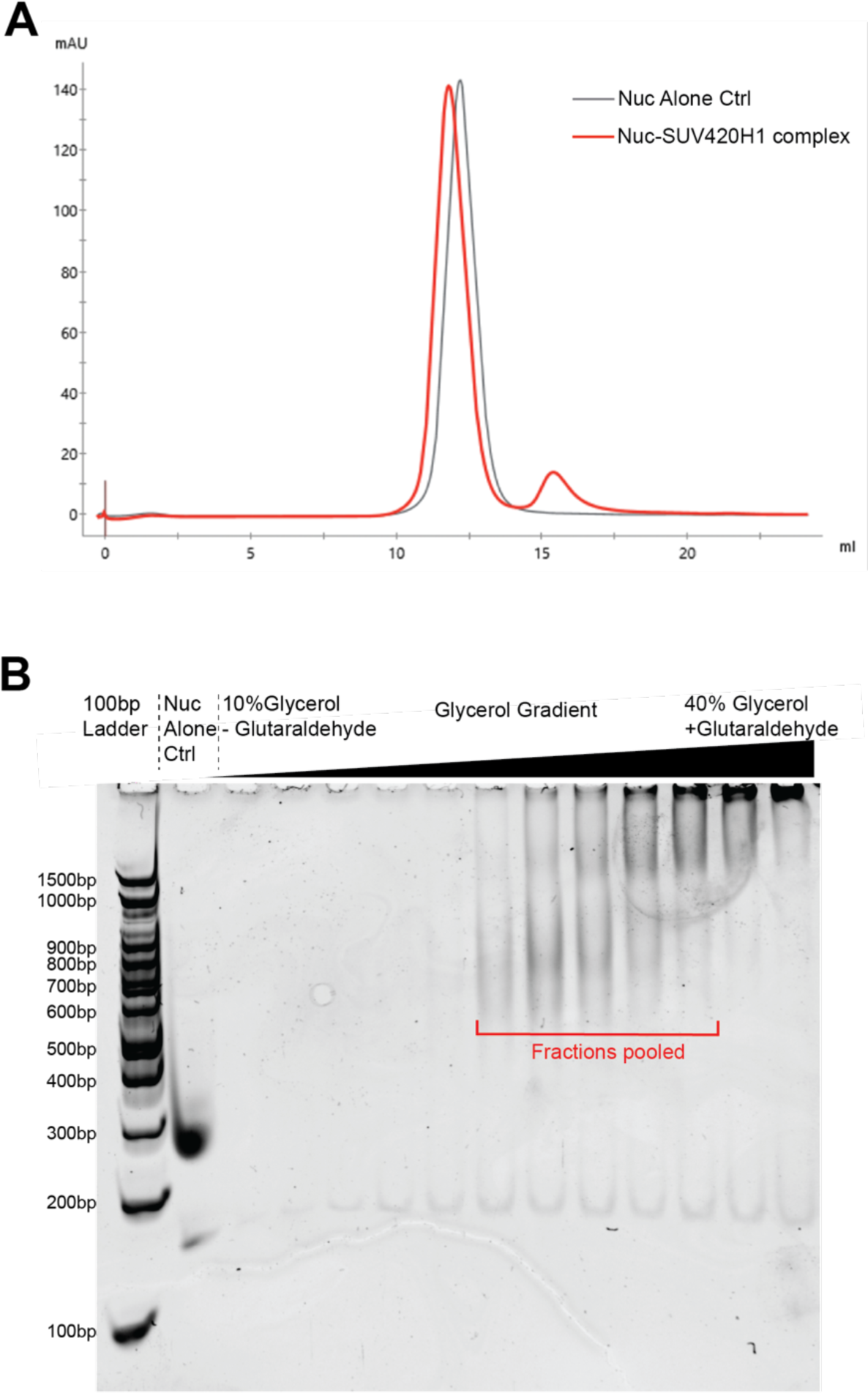
A. Gel filtration profile of nucleosome alone (grey) and SUV420H1 in complex with nucleosome (red). The gel filtration chromatography analysis shows a clear shift in the major peak when H2A.Z nucleosome is mixed with SUV420H1. B. Representative images of 4.5% Native PAGE, stained with ethidium bromide after GraFix fractionation. Fractions containing cross-linked species indicative of a protein complex were used in structural studies

**Figure S3.**
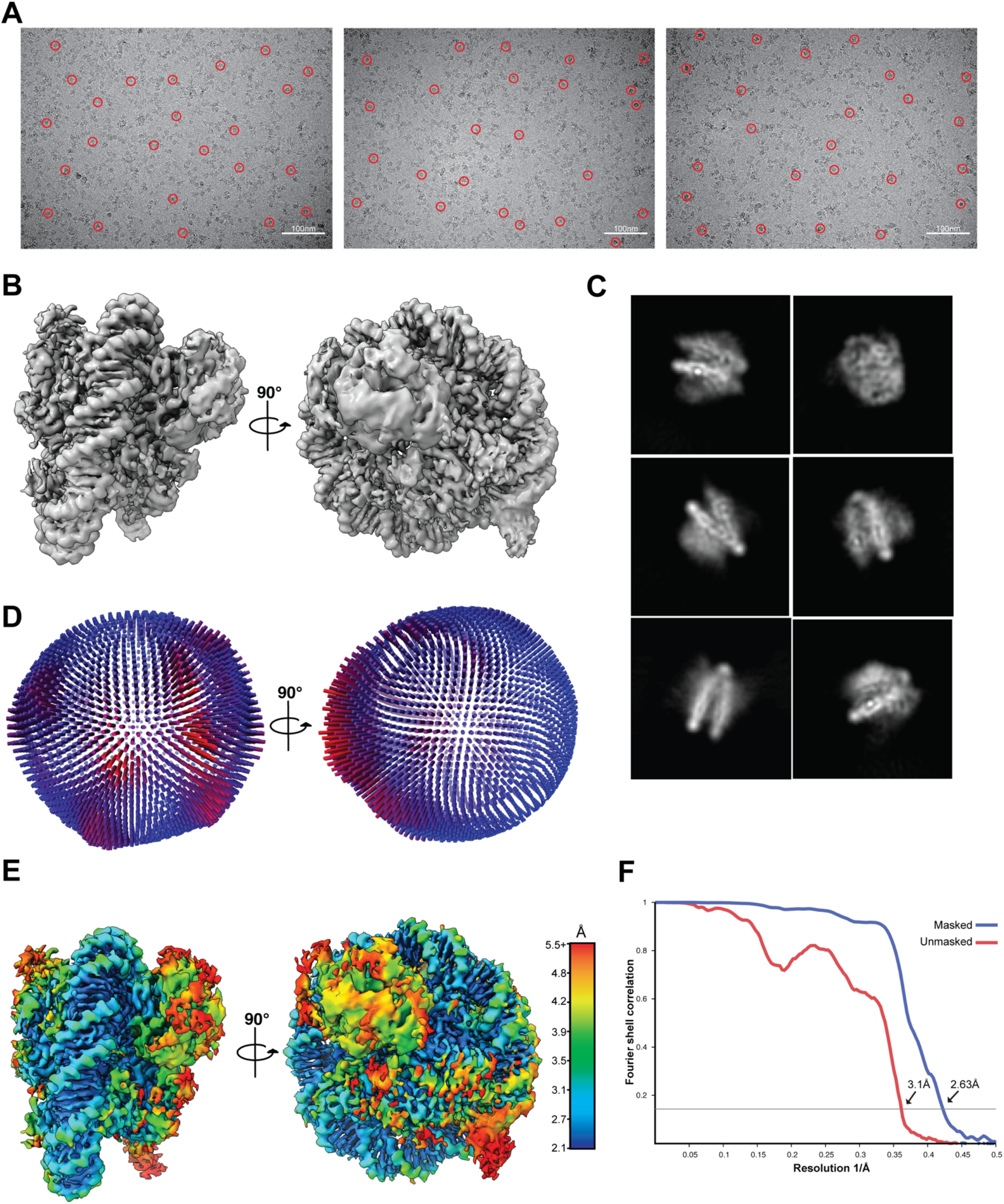
A. Representative cryo-EM micrographs of SUV420H1-H2A.Z nucleosome complex. Red circles highlight individual particles in the micrographs. B. 2.63 Å 3D reconstruction of the SUV420H1-H2A.Z nucleosome complex shown in two different views. C. Representative 2D class averages from the SUV420H1-H2A.Z nucleosome complex dataset. D. Euler angle distribution of assignment of particles used to generate the final SUV420H1-H2A.Z nucleosome complex 3D reconstruction. Cylindrical length is proportional to the number of particles assigned to the specific orientation. E. Local resolution estimation of the cryo-EM reconstructions of SUV420H1-H2A.Z nucleosome complex. F. Fourier Shell Correlation plot of the 2.63Å reconstruction of SUV420H1-H2A.Z nucleosome complex measured at FSC of 0.143.

**Figure S4.**
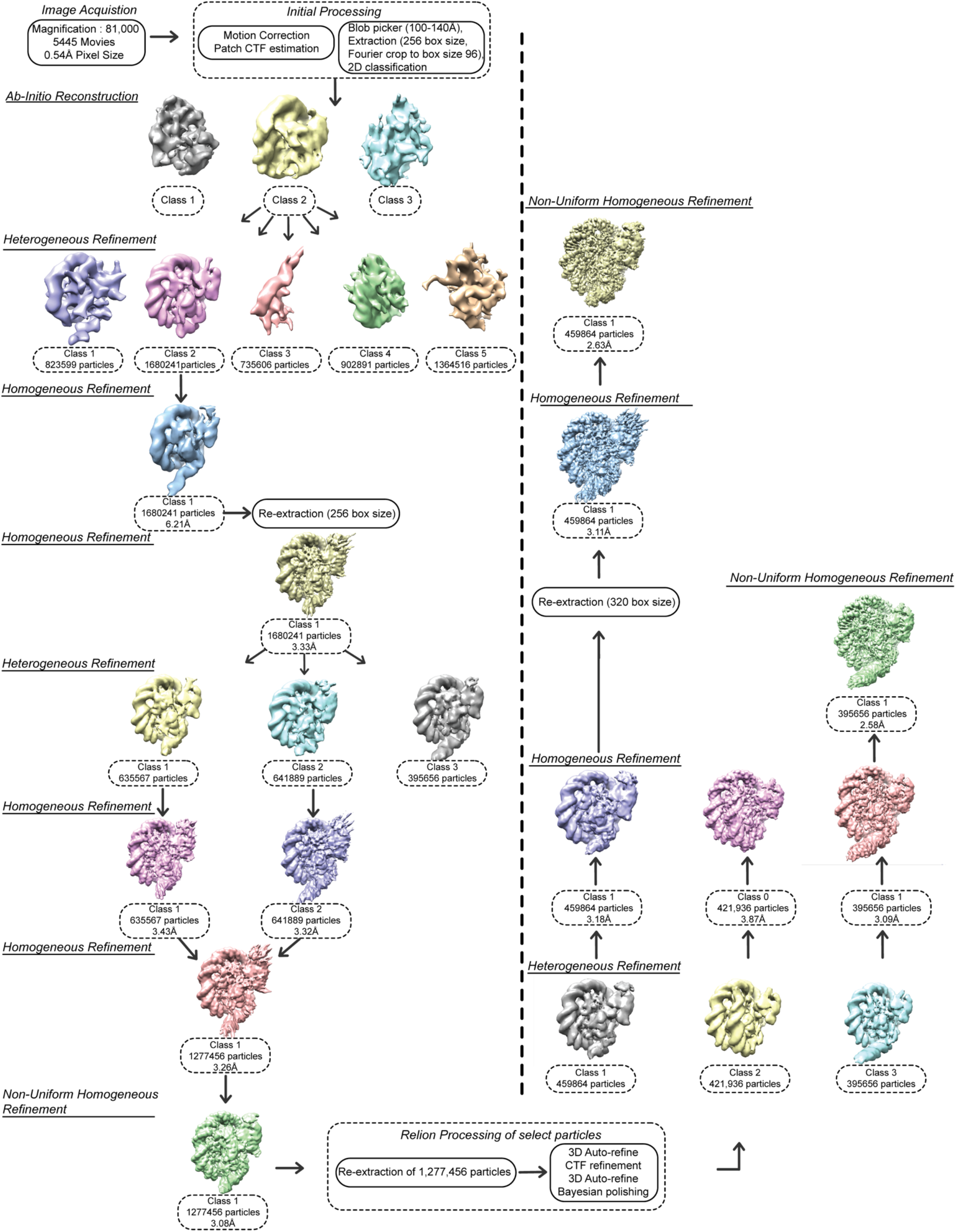
Summary of cryo-EM data processing workflow performed in cryoSPARC and RELION to obtain the final 3D reconstruction of SUV420H1-H2A.Z nucleosome complex.

**Figure S5.**
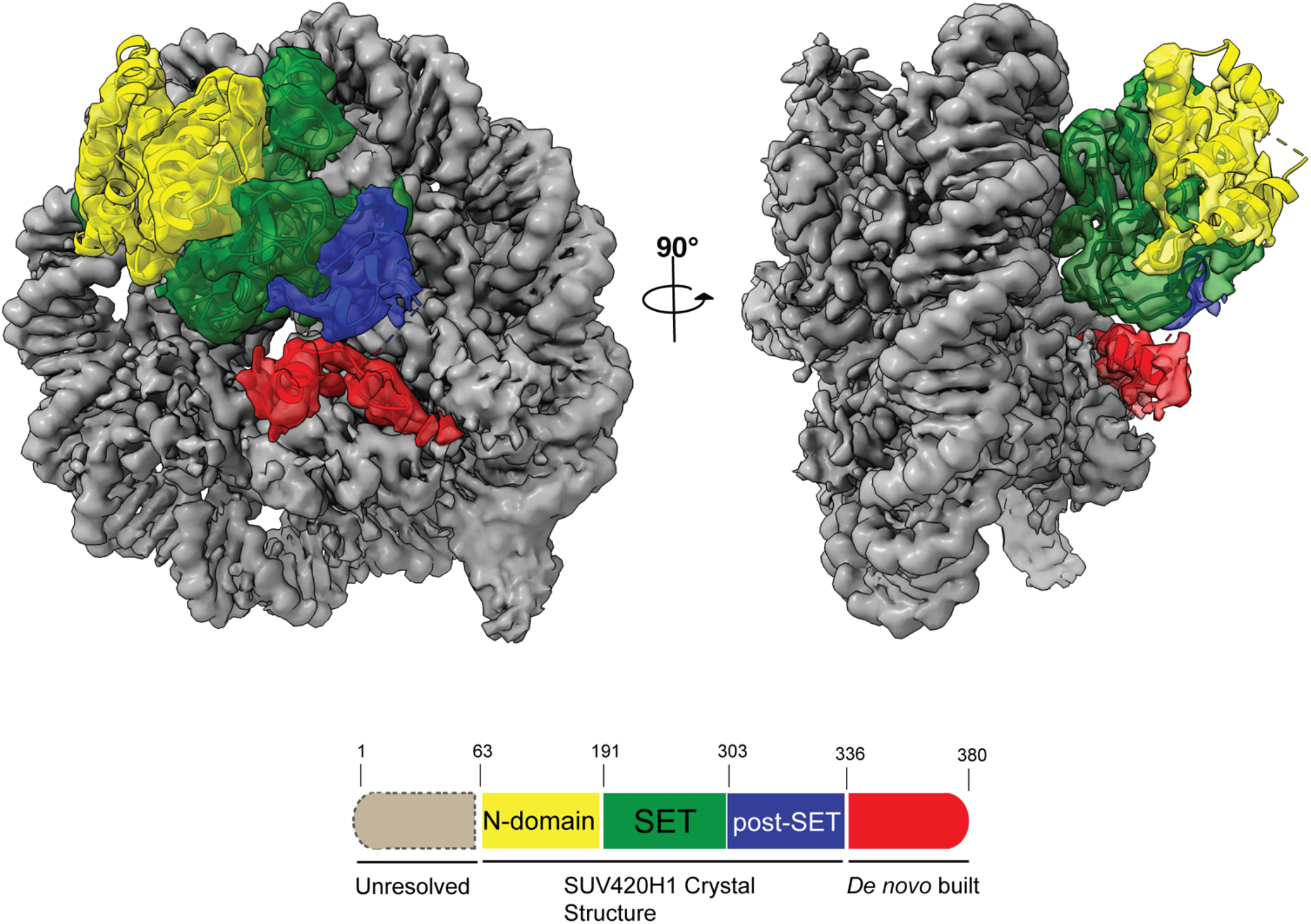
3D reconstruction SUV420H1-H2A.Z nucleosome complex colored according to the domains of SUV420H1 shown in two different views. The SUV420H1 is color-coded according to the schematic illustration below and nucleosome is colored grey. The diagram of SUV420H1, colored by domain organization, which highlights density observed and unresolved from the cryo-EM map as well as region previously determined by X-ray crystallography (PDB:3S8P).

**Figure S6.**
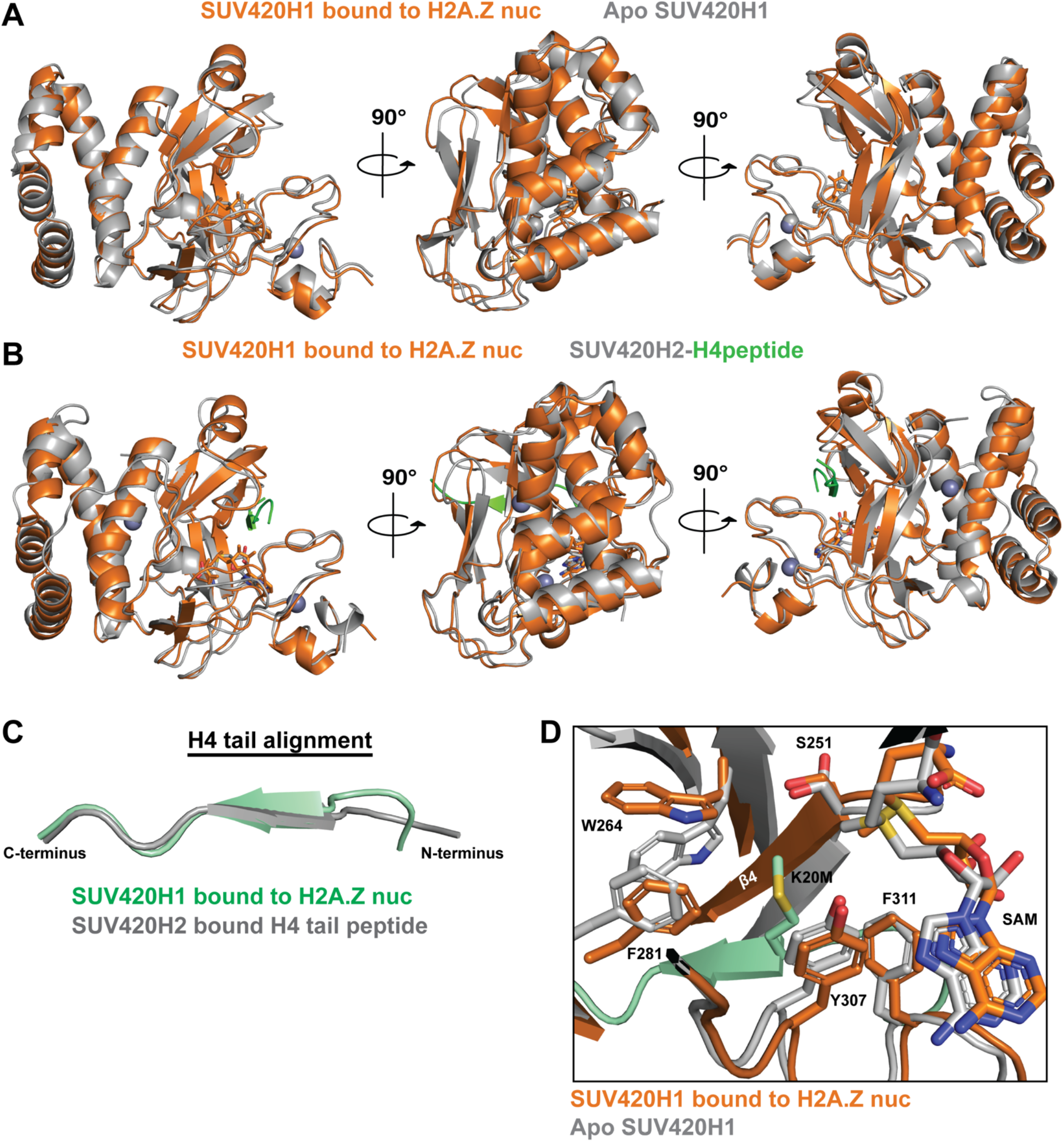
A. Structural comparison of SUV420H1-H2A.Z nucleosome complex (orange) with the Apo SUV420H1 (grey) (PDB:3S8P). The structure of SUV420H1 on the nucleosome and the apo structure containing SAM alone had a root mean square deviation (RMSD) of 0.94Å over 268 residues. B. Structural alignment of SUV420H1-H2A.Z nucleosome complex (orange) with the SUV420H2-H4 peptide complex (grey with H4 peptide in green) (PDB:4AU7). The structure of SUV420H1 on the nucleosome and the peptide bound SUV420H2 had a RMSD of 1.03Å over 268 residues C. Alignment of histone H4 tail from SUV420H1-H2A.Z nucleosome complex with H4 peptide from SUV420H2-H4 peptide complex (PDB:4AU7). The tails have an RMSD of 0.715Å over H4 residues 17-25. D. Superposition of SUV420H1-H2A.Z nucleosome complex (orange with H4 tail in green) with apo SUV420H1 (grey) showing catalytic and aromatic cage residues in SET domain.

**Figure S7.**
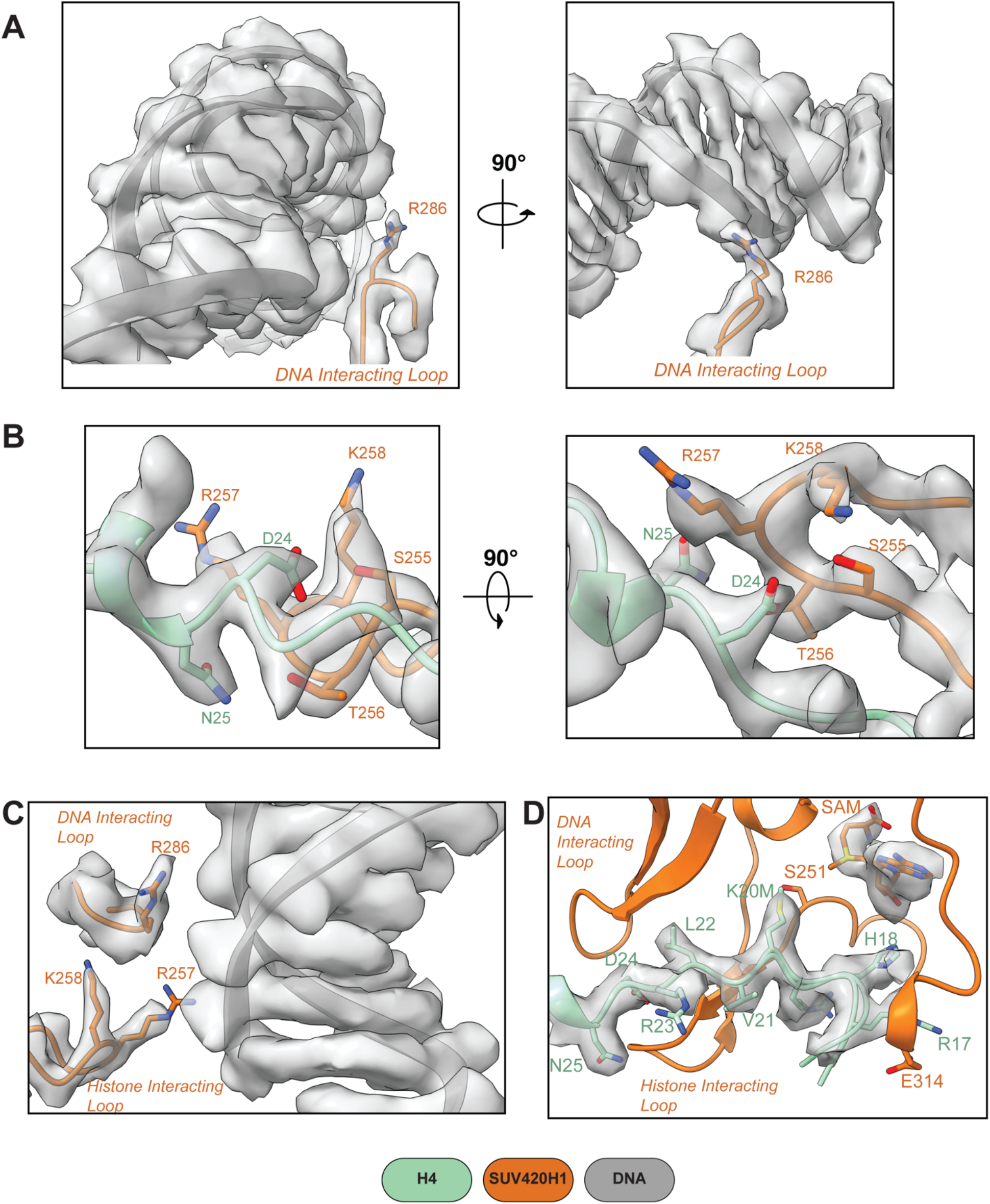
A. Cryo-EM density fit for SUV420H1 DNA interacting loop residue R286 shown in two different views. B. Density fit for SUV420H1 histone-interacting loop residues and histone H4 shown in two different views. C. Cryo-EM density views of the DNA-interacting loop, histone-interacting loop, and DNA. D. Cryo-EM density fit showing detailed interactions of histone H4 in the SET domain of SUV420H1.

**Figure S8.**
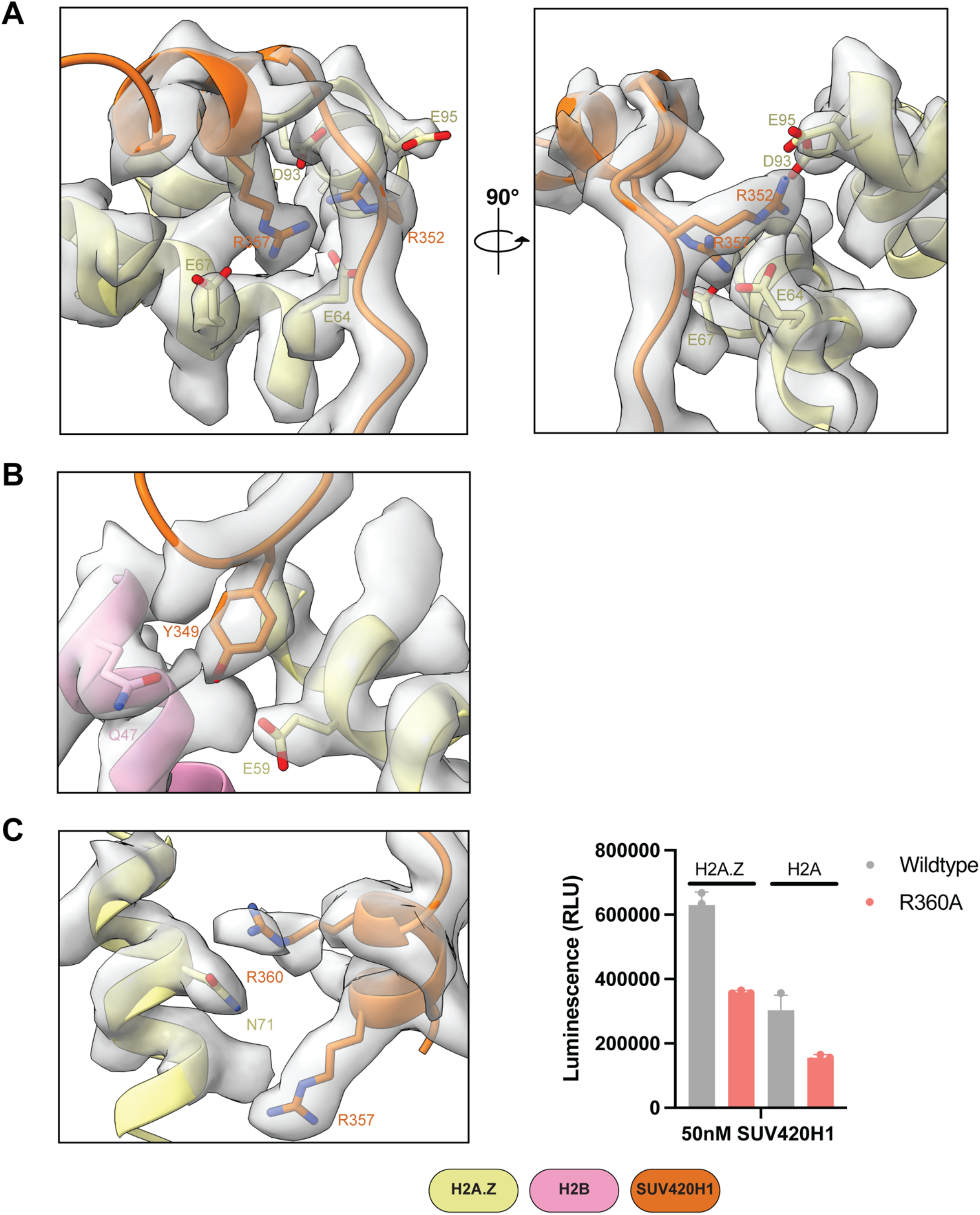
A. Close-up of the contacts between SUV420H1 and acidic patch of the nucleosome with cryo-EM density fit in two different views. B. Local cryo-EM density of the SUV420H1 residues below the canonical acidic patch interacting region. C. Local cryo-EM density of the SUV420H1 residues above the canonical acidic patch interacting region with a representative methyltransferase activity of wildtype SUV420H1 and R360A mutant on H2A and H2A.Z nucleosome.

**Figure S9.**
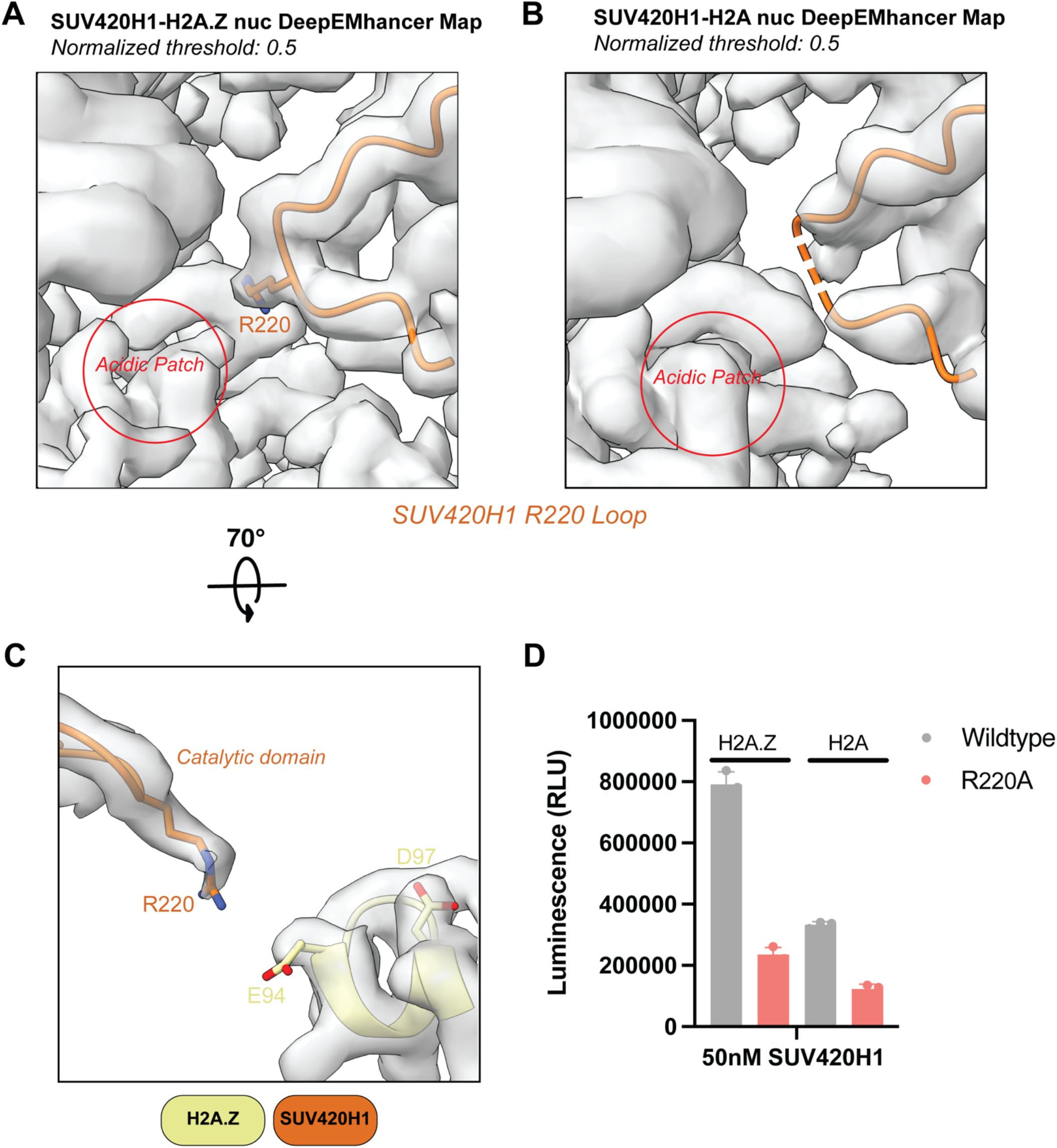
A. Cryo-EM density fit of SUV420H1 loop with R220 oriented towards the H2A.Z nucleosome acidic patch B. Cryo-EM density showing no density for loop containing R220 in H2A nucleosome structure C. Cryo-EM density fit of SUV420H1 R220 and its proximity to nucleosome acidic patch residues. D. Methyltransferase activity of wildtype SUV420H1 and R220A mutant on H2A and H2A.Z nucleosome. The data points and error bars for the methyltransferase assay represents the mean ± SD from three independent experiments.

**Figure S10.**
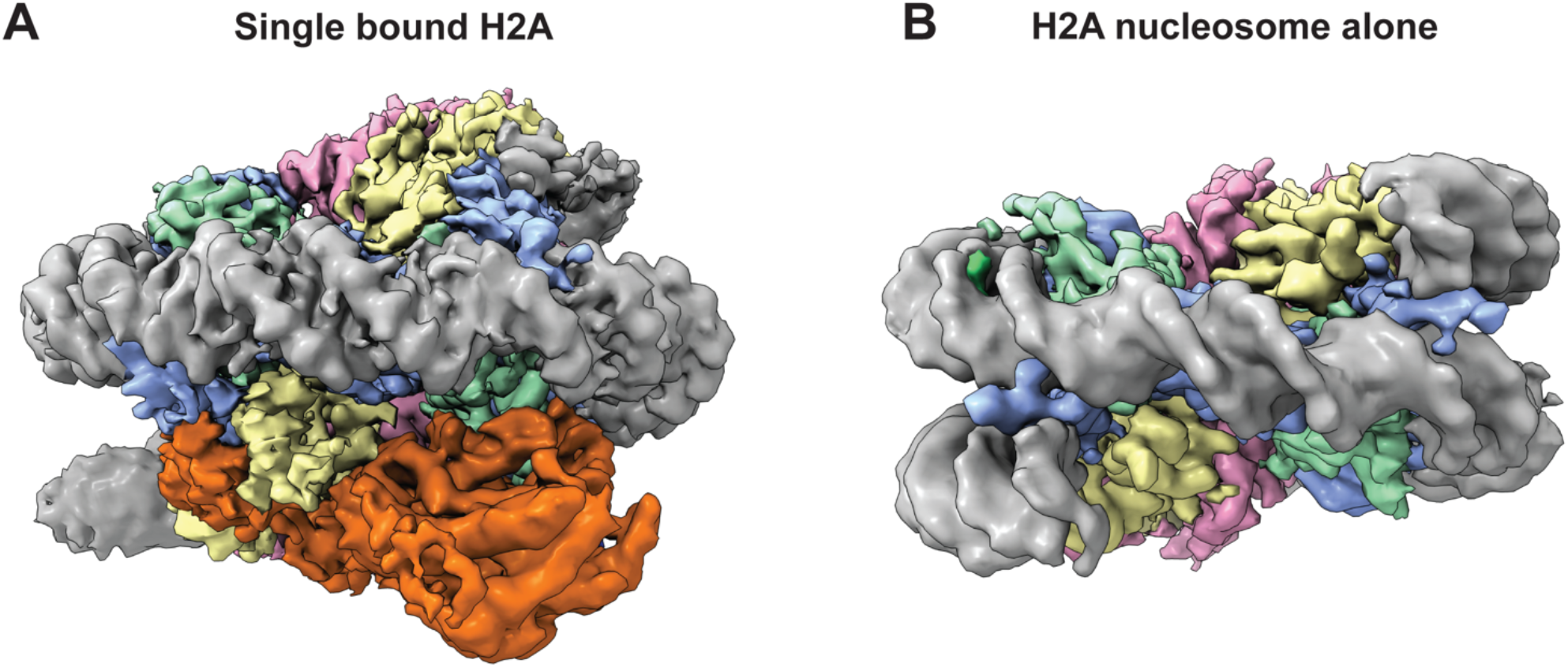
A. Cryo-EM reconstruction of SUV420H1 bound to one face of the H2A nucleosome. B. Cryo-EM reconstruction of H2A nucleosome alone.

**Figure S11.**
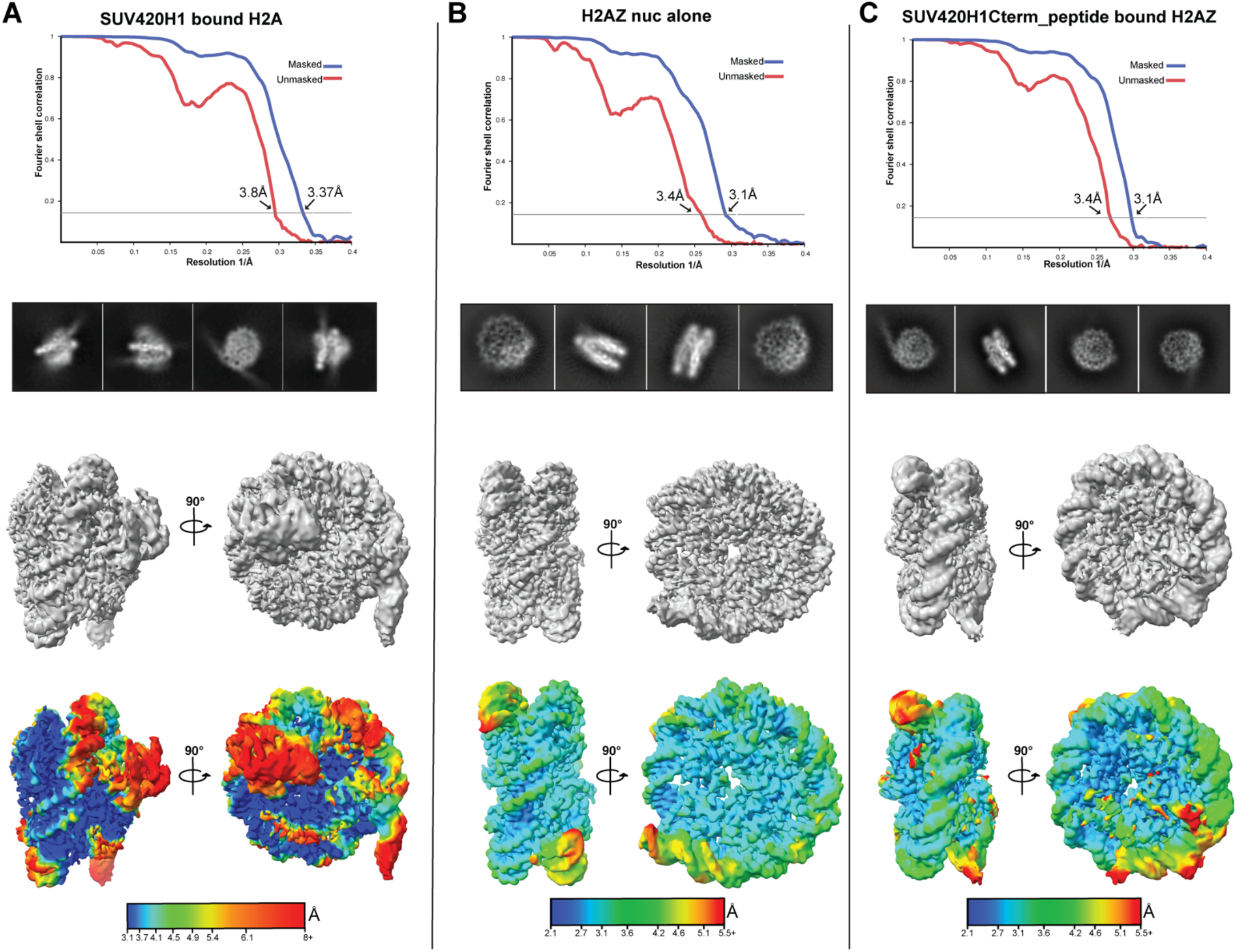
Fourier Shell Correlation plot measured at FSC of 0.143, representative 2D class averages, 3D reconstruction and local resolution estimation shown in two different views of (A) SUV420H1-H2A nucleosome complex (B) H2A.Z/H4K20M nucleosome alone and (C) SUV420H1 C-terminal peptide bound to H2A.Z nucleosome.

**Figure S12.**
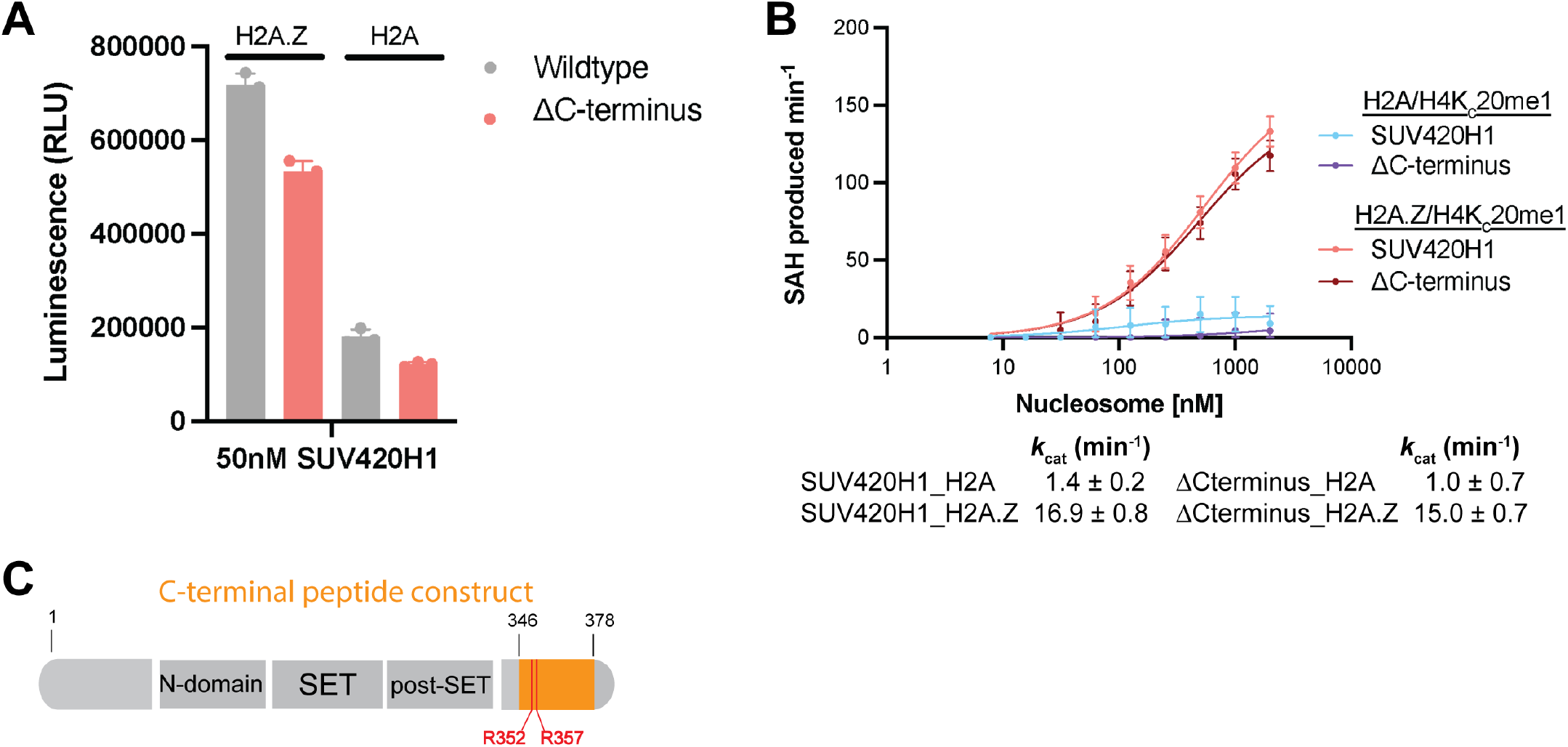
A. Representative methyltransferase activity of wildtype SUV420H1 and ΔC-terminus mutant on H2A/H4K_C_20me1 and H2A.Z/H4K_C_20me1 nucleosome. The data points and error bars for the methyltransferase assay represents the mean ± SD from three independent experiments. B. Michaelis–Menten titrations of H4K_C_20me1 nucleosomes containing H2A and H2A.Z with SUV420H1 and ΔC- terminus mutant with kinetic values quantified below. The *K*_M_ and *k*_cat_ values of the fitted data are reported in the graph. Each data point and error bar represent the mean ± SD from three independent experiments. C. Diagram of SUV420H1 highlighting the region used to synthesize the SUV420H1 C-terminal peptide construct. The two acidic patch interacting residues R352 and R357 are displayed in the diagram.

**Figure S13.**
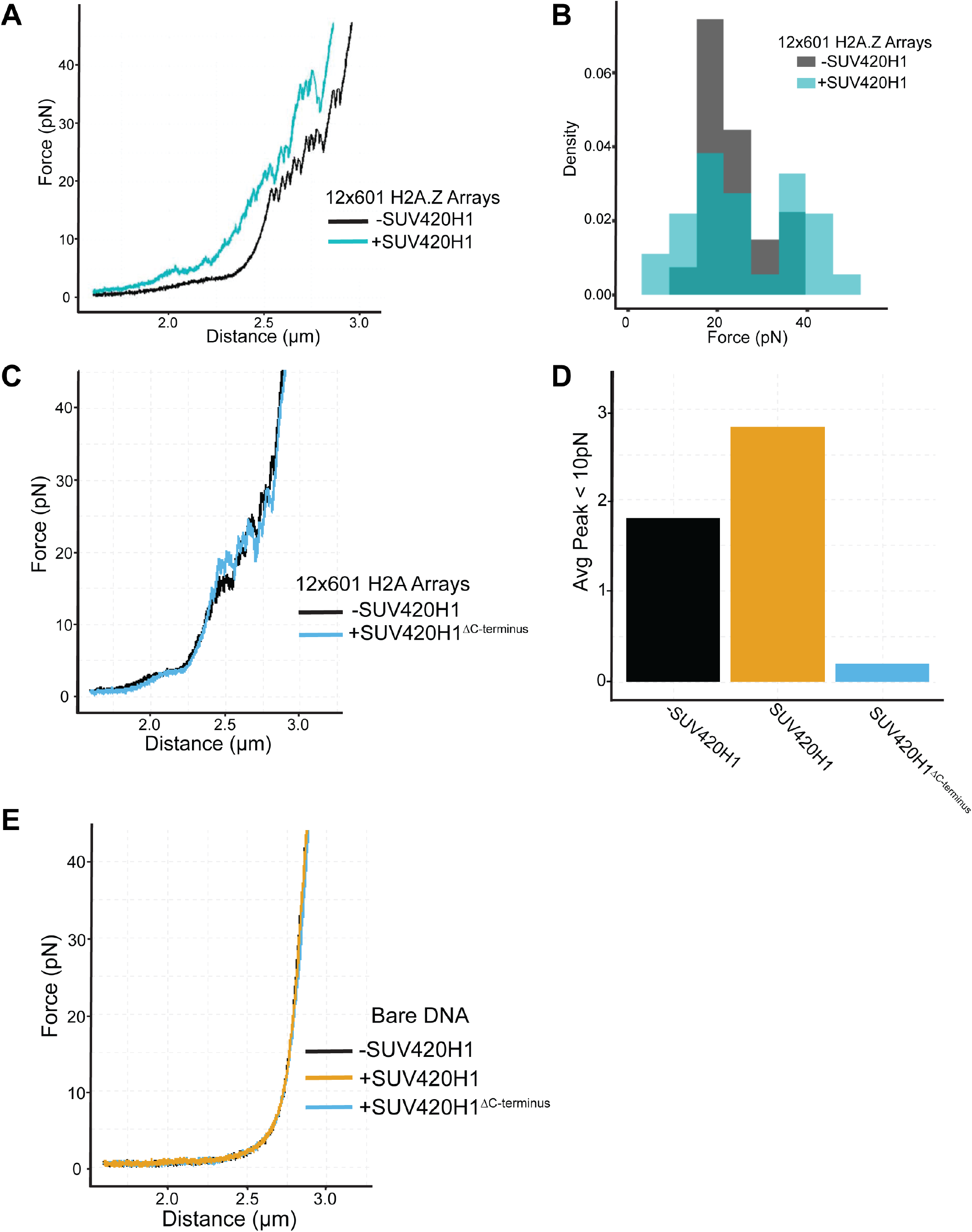
A. A representative force-extension curve for an unbound histone H2A.Z 12-mer array (black) and one for a SUV420H1- bound array (cyan). B. Distribution of transition forces observed in the force-extension curve in (A). C. A representative force-extension curve for an unbound histone H2A 12-mer array (black) and one for a ΔC-terminus SUV420H1 -bound array (blue). D. Observed average number of peaks present at less than 10pN in force-extension curve of unbound array, wildtype and ΔC-terminus SUV420H1 bound arrays. E. A representative force-extension curve for DNA alone (black) compared to wildtype (yellow) and ΔC-terminus (blue) SUV420H1 with bare DNA.

**Figure S14.**
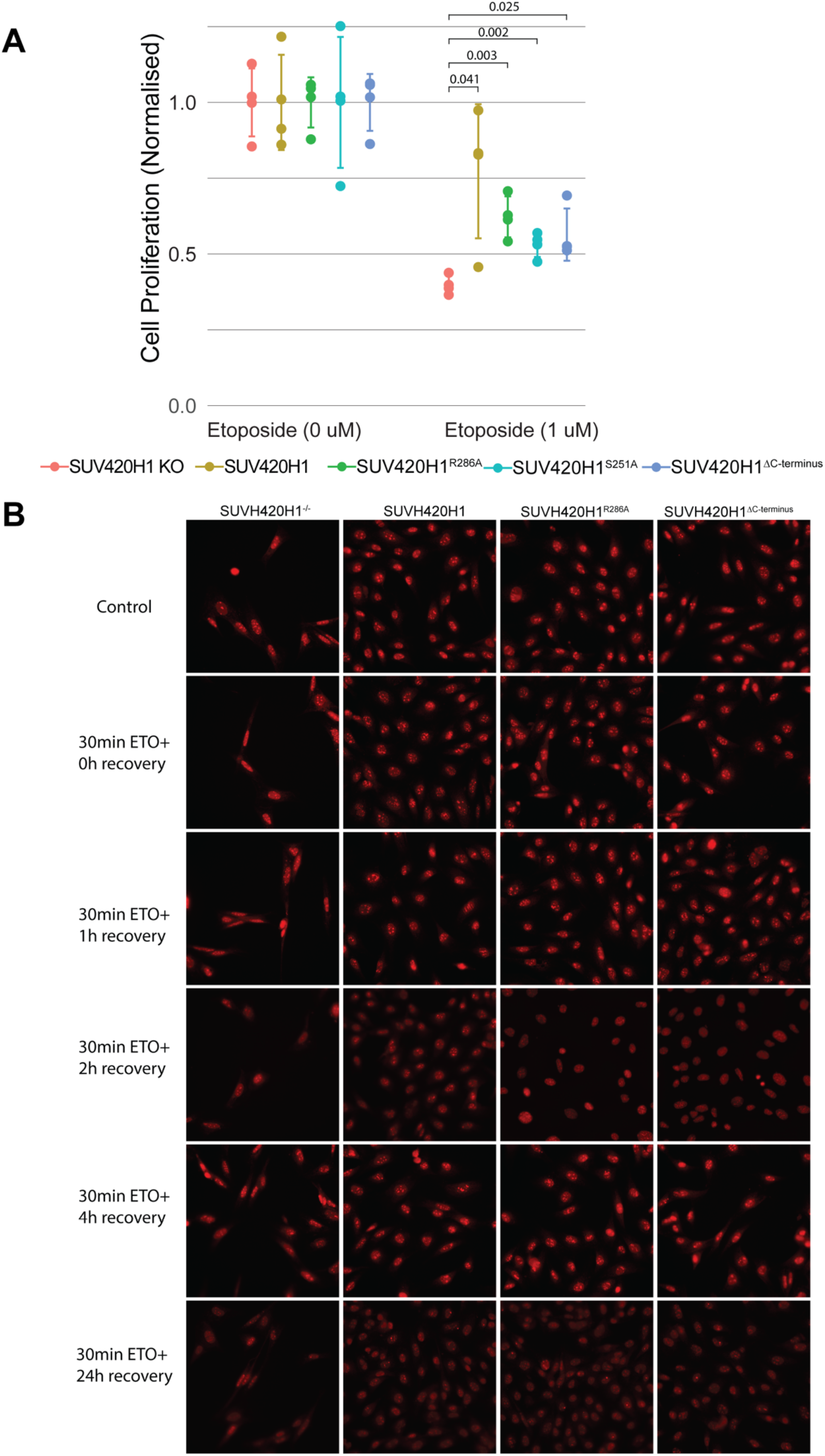
A. Cell proliferation assay to measure the sensitivity of Suv420 mutant MEFs to DNA damage induced by etoposide. Crystal violet staining was used to measure the ratio of surviving cells relative to the nontreated samples. The calculated p values from statistical analysis is presented above treated samples (4 replicates). B. Representative images of 53BP1 foci (red) staining in knockout, wildtype, R286A and ΔC-terminus SUV420H1 in MEFs cells before and after etoposide treatment.

**Table S1.** Cross-links between the SUV420H1 and H2A.Z nucleosome filtering out ones with peptide counts (PSMs) below 3.

XLMS analysis of SUV420H1/H2A.Z complex
| Protein1 | Residue | Protein2 | Residue | PSMs | Protein1 | Residue | Protein2 | Residue | PSMs | Protein1 | Residue | Protein2 | Residue | PSMs |
| --- | --- | --- | --- | --- | --- | --- | --- | --- | --- | --- | --- | --- | --- | --- |
| H2B | 109 | H3.2 | 5 | 18 | H2A.Z | 116 | H3.2 | 57 | 5 | H2B | 117 | H2A.Z | 8 | 3 |
| SUV420H1 | 116 | H3.2 | 5 | 17 | H4 | 32 | H3.2 | 28 | 5 | H2B | 117 | H2A.Z | 16 | 3 |
| H2A.Z | 116 | H3.2 | 5 | 14 | SUV420H1 | 139 | H3.2 | 5 | 5 | SUV420H1 | 116 | H2B | 47 | 3 |
| SUV420H1 | 349 | H2B | 47 | 13 | H2B | 117 | H3.2 | 28 | 5 | SUV420H1 | 171 | H3.2 | 5 | 3 |
| H2B | 109 | SUV420H1 | 364 | 12 | SUV420H1 | 171 | H3.2 | 19 | 5 | SUV420H1 | 281 | H4 | 13 | 3 |
| H4 | 32 | H3.2 | 5 | 11 | H3.2 | 24 | H2A.Z | 126 | 5 | H3.2 | 24 | SUV420H1 | 45 | 3 |
| H2A.Z | 102 | H4 | 92 | 11 | H3.2 | 15 | SUV420H1 | 173 | 5 | H3.2 | 28 | SUV420H1 | 45 | 3 |
| SUV420H1 | 173 | H3.2 | 19 | 11 | H2B | 109 | H3.2 | 28 | 5 | H3.2 | 28 | H2A.Z | 78 | 3 |
| SUV420H1 | 139 | H3.2 | 5 | 11 | SUV420H1 | 281 | H2B | 109 | 5 | H3.2 | 15 | SUV420H1 | 45 | 3 |
| H2A.Z | 75 | SUV420H1 | 364 | 11 | SUV420H1 | 389 | H2A.Z | 16 | 5 | SUV420H1 | 180 | H3.2 | 19 | 3 |
| H4 | 80 | H3.2 | 80 | 10 | SUV420H1 | 389 | H3.2 | 5 | 5 | H2B | 109 | H2A | 96 | 3 |
| H2B | 109 | H3.2 | 80 | 10 | H3.2 | 57 | H2A.Z | 126 | 5 | H2B | 109 | SUV420H1 | 334 | 3 |
| H2B | 109 | SUV420H1 | 116 | 10 | SUV420H1 | 349 | H2A.Z | 8 | 5 | SUV420H1 | 389 | H2A.Z | 8 | 3 |
| SUV420H1 | 21 | H3.2 | 28 | 10 | SUV420H1 | 349 | H2A.Z | 16 | 5 | SUV420H1 | 389 | H3.2 | 19 | 3 |
| SUV420H1 | 349 | H2B | 117 | 10 | H2B | 47 | H3.2 | 28 | 5 | SUV420H1 | 389 | H3.2 | 24 | 3 |
| SUV420H1 | 370 | H3.2 | 5 | 9 | SUV420H1 | 9 | H3.2 | 28 | 5 | SUV420H1 | 389 | H3.2 | 57 | 3 |
| H4 | 92 | H2B | 86 | 9 | H3.2 | 57 | H2A.Z | 78 | 5 | H3.2 | 24 | SUV420H1 | 45 | 3 |
| SUV420H1 | 370 | H2A.Z | 78 | 9 | SUV420H1 | 156 | H3.2 | 19 | 4 | H3.2 | 123 | H2A.Z | 126 | 3 |
| SUV420H1 | 370 | H2B | 109 | 9 | SUV420H1 | 293 | H3.2 | 19 | 4 | SUV420H1 | 349 | H2B | 6 | 3 |
| H2B | 109 | H4 | 9 | 9 | SUV420H1 | 146 | H3.2 | 19 | 4 | H2B | 47 | H3.2 | 19 | 3 |
| H2A.Z | 116 | H3.2 | 123 | 8 | H2B | 117 | H3.2 | 5 | 4 | H2B | 47 | H4 | 92 | 3 |
| SUV420H1 | 293 | H3.2 | 5 | 8 | SUV420H1 | 116 | H3.2 | 28 | 4 |  |  |  |  |  |
| SUV420H1 | 364 | H2A.Z | 78 | 8 | SUV420H1 | 217 | H3.2 | 5 | 4 |  |  |  |  |  |
| SUV420H1 | 370 | H3.2 | 19 | 8 | H3.2 | 24 | SUV420H1 | 177 | 4 |  |  |  |  |  |
| SUV420H1 | 21 | H3.2 | 5 | 8 | H3.2 | 28 | H2A.Z | 126 | 4 |  |  |  |  |  |
| H3.2 | 57 | H2A.Z | 78 | 8 | H3.2 | 28 | SUV420H1 | 177 | 4 |  |  |  |  |  |
| H4 | 92 | H2B | 86 | 8 | SUV420H1 | 370 | H3.2 | 5 | 4 |  |  |  |  |  |
| H2A.Z | 116 | H3.2 | 123 | 7 | H2B | 109 | SUV420H1 | 349 | 4 |  |  |  |  |  |
| SUV420H1 | 146 | H3.2 | 5 | 7 | H2B | 109 | SUV420H1 | 9 | 4 |  |  |  |  |  |
| SUV420H1 | 116 | H3.2 | 19 | 7 | SUV420H1 | 177 | H3.2 | 19 | 4 |  |  |  |  |  |
| H2A.Z | 126 | H3.2 | 19 | 7 | SUV420H1 | 21 | H3.2 | 19 | 4 |  |  |  |  |  |
| H4 | 80 | H3.2 | 80 | 7 | SUV420H1 | 389 | H3.2 | 28 | 4 |  |  |  |  |  |
| SUV420H1 | 370 | H3.2 | 5 | 7 | SUV420H1 | 394 | H3.2 | 19 | 4 |  |  |  |  |  |
| H2B | 109 | H4 | 13 | 7 | SUV420H1 | 394 | H2A.Z | 16 | 4 |  |  |  |  |  |
| H2B | 109 | H3.2 | 19 | 7 | H2B | 6 | H2A.Z | 8 | 4 |  |  |  |  |  |
| H2B | 6 | H3.2 | 5 | 7 | H3.2 | 123 | SUV420H1 | 293 | 4 |  |  |  |  |  |
| SUV420H1 | 349 | H3.2 | 28 | 7 | H4 | 92 | H3.2 | 57 | 4 |  |  |  |  |  |
| H2B | 47 | H3.2 | 5 | 7 | SUV420H1 | 349 | H3.2 | 57 | 4 |  |  |  |  |  |
| SUV420H1 | 9 | H3.2 | 5 | 7 | H2B | 47 | H2A.Z | 8 | 4 |  |  |  |  |  |
| SUV420H1 | 156 | H3.2 | 5 | 6 | H2B | 47 | H2A.Z | 16 | 4 |  |  |  |  |  |
| H2A.Z | 116 | SUV420H1 | 45 | 6 | SUV420H1 | 9 | H3.2 | 19 | 4 |  |  |  |  |  |
| H3.2 | 28 | SUV420H1 | 293 | 6 | SUV420H1 | 212 | H3.2 | 15 | 4 |  |  |  |  |  |
| SUV420H1 | 180 | H3.2 | 5 | 6 | SUV420H1 | 45 | H3.2 | 19 | 3 |  |  |  |  |  |
| SUV420H1 | 370 | H3.2 | 57 | 6 | H2A.Z | 116 | H3.2 | 5 | 3 |  |  |  |  |  |
| SUV420H1 | 349 | H3.2 | 19 | 6 | H2B | 121 | H2A.Z | 16 | 3 |  |  |  |  |  |
| H3.2 | 57 | H2A.Z | 126 | 6 | H4 | 32 | H3.2 | 65 | 3 |  |  |  |  |  |
| SUV420H1 | 212 | H3.2 | 5 | 6 | SUV420H1 | 153 | H3.2 | 5 | 3 |  |  |  |  |  |
| SUV420H1 | 45 | H3.2 | 5 | 5 | H4 | 60 | SUV420H1 | 293 | 3 |  |  |  |  |  |

**Table S2.** Cryo-EM data collection, refinement and validation statistics

Cryo-EM data collection, refinement and validation statistics
|  | #1 SUV420H1-H2A.Z<br>(EMDB-xxxx)<br>(PDB xxxx) | #2 SUV420H1-H2A<br>(EMDB-xxxx)<br>(PDB xxxx) | #3 H2A.Z nuc<br>(EMDB-xxxx)<br>(PDB xxxx) | #4 SUV420H1 C-terminus<br>peptide-H2A.Z<br>nuc<br>(EMDB-xxxx)<br>(PDB xxxx) | #5 H2A nuc |
| --- | --- | --- | --- | --- | --- |
| **Data collection and processing** |  |  |  |  |  |
| Magnification | 81,000 | 81,000 | 105,000 | 36,000 | 36,000 |
| Voltage (kV) | 300 | 300 | 300 | 200 | 200 |
| Electron exposure (e-/Å<sup>2</sup>) | 50 | 52.1 | 54.76 | 50.89 | 50.89 |
| Defocus range (µm) | -1 to -2.4 | -0.25 to -1.75 | -1.3 to -1.5 | -1.0 to -2.9 | -1.0 to -2.9 |
| Pixel size (Å) | 1.08 | 1.08 | 0.84 | 1.096 | 1.096 |
| Symmetry imposed | C1 | C1 | C1 | C1 | C1 |
| Initial particle images (no.) | 5,506,853 | 4,761,686 | 962,712 | 1,615,457 | 220,145 |
| Final particle images (no.) | 459,864 | 366,390 | 206,872 | 494,524 | 79,392 |
| Map resolution (Å)<br>FSC threshold 0.143 | 2.63 | 3.37 | 3.1 | 3.1 | 3.86 |
| Map resolution range (Å) | 2.4-27.9 | 2.7-51.6 | 2.6-49.5 | 2.5-46.2 | 2.4-8.7 |
| **Refinement** |  |  |  |  |  |
| Initial model used (PDB code) | 3S8P, 1F66, 3TU4 | 3S8P, 1AOI, 3TU4 | 1F66 | SUV420H1-H2A.Z complex | * |
| Map sharpening *B* factor (Å<sup>2</sup>) | -85.2 | -122.0 | -87.2 | -105.9 | -175.8 |
| Model composition |  |  |  |  | * |
| Non-hydrogen atoms | 13,183 | 13,300 | 12,063 | 12,166 |  |
| Protein residues | 1,034 | 1,046 | 769 | 783 |  |
| Ligands | 2 | 2 | 0 | 0 |  |
| *B* factors (Å<sup>2</sup>) |  |  |  |  | * |
| Protein | 70.18 | 41.63 | 26.98 | 118.09 |  |
| Ligand | 116.50 | 110.28 | 84.25 | 234.14 |  |
| R.m.s. deviations |  |  |  |  | * |
| Bond lengths (Å) | 0.008 | 0.006 | 0.007 | 0.007 |  |
| Bond angles (°) | 0.849 | 0.626 | 0.845 | 0.698 |  |
| Validation |  |  |  |  | * |
| MolProbity score | 1.24 | 1.81 | 2.09 | 2.50 |  |
| Clashscore | 2.88 | 6.32 | 3.09 | 9.05 |  |
| Poor rotamers (%) | 1.16 | 0.00 | 7.54 | 6.78 |  |
| Ramachandran plot |  |  |  |  | * |
| Favored (%) | 97.43 | 92.58 | 95.35 | 94.38 |  |
| Allowed (%) | 2.57 | 7.42 | 4.65 | 5.62 |  |
| Disallowed (%) | 0.00 | 0.00 | 0.00 | 0.00 |  |

\* Model was not built

## References

Adams, P.D., Afonine, P.V., Bunkoczi, G., Chen, V.B., Davis, I.W., Echols, N., Headd, J.J., Hung, L.W., Kapral, G.J., Grosse-Kunstleve, R.W., et al. (2010). PHENIX: a comprehensive Python-based system for macromolecular structure solution. Acta Crystallogr D Biol Crystallogr 66, 213–221.

Armache, K.J., Garlick, J.D., Canzio, D., Narlikar, G.J., and Kingston, R.E. (2011). Structural basis of silencing: Sir3 BAH domain in complex with a nucleosome at 3.0 A resolution. Science 334, 977–982.

Brohm, A., Elsawy, H., Rathert, P., Kudithipudi, S., Schoch, T., Schuhmacher, M.K., Weirich, S., and Jeltsch, A. (2019). Somatic Cancer Mutations in the SUV420H1 Protein Lysine Methyltransferase Modulate Its Catalytic Activity. J Mol Biol 431, 3068–3080.

Bromberg, K.D., Mitchell, T.R., Upadhyay, A.K., Jakob, C.G., Jhala, M.A., Comess, K.M., Lasko, L.M., Li, C., Tuzon, C.T., Dai, Y., et al. (2017). The SUV4-20 inhibitor A-196 verifies a role for epigenetics in genomic integrity. Nat Chem Biol 13, 317–324.

Brower-Toland, B.D., Smith, C.L., Yeh, R.C., Lis, J.T., Peterson, C.L., and Wang, M.D. (2002). Mechanical disruption of individual nucleosomes reveals a reversible multistage release of DNA. Proc Natl Acad Sci U S A 99, 1960–1965.

Buck, S., Pekarek, L., and Caliskan, N. (2021). POTATO: An automated pipeline for batch analysis of optical tweezers data. bioRxiv.

Bustamante, C.J., Chemla, Y.R., Liu, S., and Wang, M.D. (2021). Optical tweezers in single-molecule biophysics. Nat Rev Methods Primers 1.

Cerami, E., Gao, J., Dogrusoz, U., Gross, B.E., Sumer, S.O., Aksoy, B.A., Jacobsen, A., Byrne, C.J., Heuer, M.L., Larsson, E., et al. (2012). The cBio cancer genomics portal: an open platform for exploring multidimensional cancer genomics data. Cancer Discov 2, 401–404.

Chen, Z.L., Meng, J.M., Cao, Y., Yin, J.L., Fang, R.Q., Fan, S.B., Liu, C., Zeng, W.F., Ding, Y.H., Tan, D., et al. (2019). A high-speed search engine pLink 2 with systematic evaluation for proteome-scale identification of cross-linked peptides. Nat Commun 10, 3404.

Dawson, M.A., Bannister, A.J., Gottgens, B., Foster, S.D., Bartke, T., Green, A.R., and Kouzarides, T. (2009). JAK2 phosphorylates histone H3Y41 and excludes HP1alpha from chromatin. Nature 461, 819–822.

Dialynas, G.K., Makatsori, D., Kourmouli, N., Theodoropoulos, P.A., McLean, K., Terjung, S., Singh, P.B., and Georgatos, S.D. (2006). Methylation-independent binding to histone H3 and cell cycle-dependent incorporation of HP1beta into heterochromatin. J Biol Chem 281, 14350–14360.

Dyer, P.N., Edayathumangalam, R.S., White, C.L., Bao, Y., Chakravarthy, S., Muthurajan, U.M., and Luger, K. (2004). Reconstitution of nucleosome core particles from recombinant histones and DNA. Methods Enzymol 375, 23–44.

Emsley, P., and Cowtan, K. (2004). Coot: model-building tools for molecular graphics. Acta Crystallogr D Biol Crystallogr 60, 2126–2132.

Farr, S.E., Woods, E.J., Joseph, J.A., Garaizar, A., and Collepardo-Guevara, R. (2021). Nucleosome plasticity is a critical element of chromatin liquid-liquid phase separation and multivalent nucleosome interactions. Nat Commun 12, 2883.

Gibson, B.A., Doolittle, L.K., Schneider, M.W.G., Jensen, L.E., Gamarra, N., Henry, L., Gerlich, D.W., Redding, S., and Rosen, M.K. (2019). Organization of Chromatin by Intrinsic and Regulated Phase Separation. Cell 179, 470–484 e421.

Grau, D., Zhang, Y., Lee, C.H., Valencia-Sanchez, M., Zhang, J., Wang, M., Holder, M., Svetlov, V., Tan, D., Nudler, E., et al. (2021). Structures of monomeric and dimeric PRC2:EZH1 reveal flexible modules involved in chromatin compaction. Nat Commun 12, 714.

Groth, A., Rocha, W., Verreault, A., and Almouzni, G. (2007). Chromatin challenges during DNA replication and repair. Cell 128, 721–733.

Hauer, M.H., and Gasser, S.M. (2017). Chromatin and nucleosome dynamics in DNA damage and repair. Genes Dev 31, 2204–2221.

Husmann, D., and Gozani, O. (2019). Histone lysine methyltransferases in biology and disease. Nat Struct Mol Biol 26, 880–889.

Illingworth, R.S., Moffat, M., Mann, A.R., Read, D., Hunter, C.J., Pradeepa, M.M., Adams, I.R., and Bickmore, W.A. (2015). The E3 ubiquitin ligase activity of RING1B is not essential for early mouse development. Genes Dev 29, 1897–1902.

Jang, S.M., Azebi, S., Soubigou, G., and Muchardt, C. (2014). DYRK1A phoshorylates histone H3 to differentially regulate the binding of HP1 isoforms and antagonize HP1-mediated transcriptional repression. EMBO Rep 15, 686–694.

Jorgensen, S., Schotta, G., and Sorensen, C.S. (2013). Histone H4 lysine 20 methylation: key player in epigenetic regulation of genomic integrity. Nucleic Acids Res 41, 2797–2806.

Kudithipudi, S., Dhayalan, A., Kebede, A.F., and Jeltsch, A. (2012). The SET8 H4K20 protein lysine methyltransferase has a long recognition sequence covering seven amino acid residues. Biochimie 94, 2212–2218.

Kuo, A.J., Song, J., Cheung, P., Ishibe-Murakami, S., Yamazoe, S., Chen, J.K., Patel, D.J., and Gozani, O. (2012). The BAH domain of ORC1 links H4K20me2 to DNA replication licensing and Meier-Gorlin syndrome. Nature 484, 115–119.

Larson, A.G., Elnatan, D., Keenen, M.M., Trnka, M.J., Johnston, J.B., Burlingame, A.L., Agard, D.A., Redding, S., and Narlikar, G.J. (2017). Liquid droplet formation by HP1alpha suggests a role for phase separation in heterochromatin. Nature 547, 236–240.

Lavigne, M., Eskeland, R., Azebi, S., Saint-Andre, V., Jang, S.M., Batsche, E., Fan, H.Y., Kingston, R.E., Imhof, A., and Muchardt, C. (2009). Interaction of HP1 and Brg1/Brm with the globular domain of histone H3 is required for HP1- mediated repression. PLoS Genet 5, e1000769.

Lee, S., Oh, S., Jeong, K., Jo, H., Choi, Y., Seo, H.D., Kim, M., Choe, J., Kwon, C.S., and Lee, D. (2018). Dot1 regulates nucleosome dynamics by its inherent histone chaperone activity in yeast. Nat Commun 9, 240.

Leicher, R., Ge, E.J., Lin, X., Reynolds, M.J., Xie, W., Walz, T., Zhang, B., Muir, T.W., and Liu, S. (2020). Single-molecule and in silico dissection of the interaction between Polycomb repressive complex 2 and chromatin. Proc Natl Acad Sci U S A 117, 30465–30475.

Leicher R, L.S. (2022). Probing the interaction between chromatin and chromatin-associated complexes with optical tweezers. In Optical Tweezers: Methods and Protocols (Humana Press).

Lewis, P.W., Muller, M.M., Koletsky, M.S., Cordero, F., Lin, S., Banaszynski, L.A., Garcia, B.A., Muir, T.W., Becher, O.J., and Allis, C.D. (2013). Inhibition of PRC2 activity by a gain-of-function H3 mutation found in pediatric glioblastoma. Science 340, 857–861.

Lewis, T.S., Sokolova, V., Jung, H., Ng, H., and Tan, D. (2021). Structural basis of chromatin regulation by histone variant H2A.Z. Nucleic Acids Res 49, 11379–11391.

Li, X., Mooney, P., Zheng, S., Booth, C.R., Braunfeld, M.B., Gubbens, S., Agard, D.A., and Cheng, Y. (2013). Electron counting and beam-induced motion correction enable near-atomic-resolution single-particle cryo-EM. Nature methods 10, 584–590.

Long, H., Zhang, L., Lv, M., Wen, Z., Zhang, W., Chen, X., Zhang, P., Li, T., Chang, L., Jin, C., et al. (2020). H2A.Z facilitates licensing and activation of early replication origins. Nature 577, 576–581.

Lowary, P.T., and Widom, J. (1998). New DNA sequence rules for high affinity binding to histone octamer and sequence-directed nucleosome positioning. J Mol Biol 276, 19–42.

Luger, K., Mader, A.W., Richmond, R.K., Sargent, D.F., and Richmond, T.J. (1997). Crystal structure of the nucleosome core particle at 2.8 A resolution. Nature 389, 251–260.

Martin, C., and Zhang, Y. (2005). The diverse functions of histone lysine methylation. Nat Rev Mol Cell Biol 6, 838–849.

McGinty, R.K., and Tan, S. (2015). Nucleosome structure and function. Chem Rev 115, 2255–2273.

Meng, H., Andresen, K., and van Noort, J. (2015). Quantitative analysis of single-molecule force spectroscopy on folded chromatin fibers. Nucleic Acids Res 43, 3578–3590.

Morgan, M.A.J., and Shilatifard, A. (2020). Reevaluating the roles of histone-modifying enzymes and their associated chromatin modifications in transcriptional regulation. Nat Genet 52, 1271–1281.

Nishioka, K., Rice, J.C., Sarma, K., Erdjument-Bromage, H., Werner, J., Wang, Y., Chuikov, S., Valenzuela, P., Tempst, P., Steward, R., et al. (2002). PR-Set7 is a nucleosome-specific methyltransferase that modifies lysine 20 of histone H4 and is associated with silent chromatin. Mol Cell 9, 1201–1213.

O’Reilly, F.J., and Rappsilber, J. (2018). Cross-linking mass spectrometry: methods and applications in structural, molecular and systems biology. Nat Struct Mol Biol 25, 1000–1008.

Paquin, K.L., and Howlett, N.G. (2018). Understanding the Histone DNA Repair Code: H4K20me2 Makes Its Mark. Mol Cancer Res 16, 1335–1345.

Paulsen, B., Velasco, S., Kedaigle, A.J., Pigoni, M., Quadrato, G., Deo, A.J., Adiconis, X., Uzquiano, A., Sartore, R., Yang, S.M., et al. (2022). Autism genes converge on asynchronous development of shared neuron classes. Nature 602, 268–273.

Pesavento, J.J., Yang, H., Kelleher, N.L., and Mizzen, C.A. (2008). Certain and progressive methylation of histone H4 at lysine 20 during the cell cycle. Mol Cell Biol 28, 468–486.

Pettersen, E.F., Goddard, T.D., Huang, C.C., Couch, G.S., Greenblatt, D.M., Meng, E.C., and Ferrin, T.E. (2004). UCSF Chimera--a visualization system for exploratory research and analysis. J Comput Chem 25, 1605–1612.

Punjani, A., Rubinstein, J.L., Fleet, D.J., and Brubaker, M.A. (2017). cryoSPARC: algorithms for rapid unsupervised cryo-EM structure determination. Nat Methods 14, 290–296.

Rosenthal, P.B., and Henderson, R. (2003). Optimal determination of particle orientation, absolute hand, and contrast loss in single-particle electron cryomicroscopy. J Mol Biol 333, 721–745.

Sanchez-Garcia, R., Gomez-Blanco, J., Cuervo, A., Carazo, J.M., Sorzano, C.O.S., and Vargas, J. (2021). DeepEMhancer: a deep learning solution for cryo-EM volume post-processing. Commun Biol 4, 874.

Sanulli, S., Trnka, M.J., Dharmarajan, V., Tibble, R.W., Pascal, B.D., Burlingame, A.L., Griffin, P.R., Gross, J.D., and Narlikar, G.J. (2019). HP1 reshapes nucleosome core to promote phase separation of heterochromatin. Nature 575, 390–394.

Schotta, G., Lachner, M., Sarma, K., Ebert, A., Sengupta, R., Reuter, G., Reinberg, D., and Jenuwein, T. (2004). A silencing pathway to induce H3-K9 and H4-K20 trimethylation at constitutive heterochromatin. Genes Dev 18, 1251–1262.

Schotta, G., Sengupta, R., Kubicek, S., Malin, S., Kauer, M., Callen, E., Celeste, A., Pagani, M., Opravil, S., De La Rosa-Velazquez, I.A., et al. (2008). A chromatin-wide transition to H4K20 monomethylation impairs genome integrity and programmed DNA rearrangements in the mouse. Genes Dev 22, 2048–2061.

Schrodinger, LLC (2015). The PyMOL Molecular Graphics System, Version 1.8.

Shan, C.M., Wang, J., Xu, K., Chen, H., Yue, J.X., Andrews, S., Moresco, J.J., Yates, J.R., Nagy, P.L., Tong, L., et al. (2016). A histone H3K9M mutation traps histone methyltransferase Clr4 to prevent heterochromatin spreading. Elife 5.

Shechter, D., Dormann, H.L., Allis, C.D., and Hake, S.B. (2007). Extraction, purification and analysis of histones. Nat Protoc 2, 1445–1457.

Simon, M.D. (2010). Installation of site-specific methylation into histones using methyl lysine analogs. Curr Protoc Mol Biol Chapter 21, Unit 21 18 21-10.

Southall, S.M., Cronin, N.B., and Wilson, J.R. (2014). A novel route to product specificity in the Suv4-20 family of histone H4K20 methyltransferases. Nucleic Acids Res 42, 661–671.

Stark, H. (2010). GraFix: stabilization of fragile macromolecular complexes for single particle cryo-EM. Methods Enzymol 481, 109–126.

Strom, A.R., Emelyanov, A.V., Mir, M., Fyodorov, D.V., Darzacq, X., and Karpen, G.H. (2017). Phase separation drives heterochromatin domain formation. Nature 547, 241–245.

Suloway, C., Pulokas, J., Fellmann, D., Cheng, A., Guerra, F., Quispe, J., Stagg, S., Potter, C.S., and Carragher, B. (2005). Automated molecular microscopy: the new Leginon system. J Struct Biol 151, 41–60.

Suto, R.K., Clarkson, M.J., Tremethick, D.J., and Luger, K. (2000). Crystal structure of a nucleosome core particle containing the variant histone H2A.Z. Nat Struct Biol 7, 1121–1124.

Sze, C.C., Cao, K., Collings, C.K., Marshall, S.A., Rendleman, E.J., Ozark, P.A., Chen, F.X., Morgan, M.A., Wang, L., and Shilatifard, A. (2017). Histone H3K4 methylation-dependent and -independent functions of Set1A/COMPASS in embryonic stem cell self-renewal and differentiation. Genes Dev 31, 1732–1737.

Taglialatela, A., Leuzzi, G., Sannino, V., Cuella-Martin, R., Huang, J.W., Wu-Baer, F., Baer, R., Costanzo, V., and Ciccia, A. (2021). REV1-Polzeta maintains the viability of homologous recombination-deficient cancer cells through mutagenic repair of PRIMPOL-dependent ssDNA gaps. Mol Cell 81, 4008–4025 e4007.

Tsang, L.W., Hu, N., and Underhill, D.A. (2010). Comparative analyses of SUV420H1 isoforms and SUV420H2 reveal differences in their cellular localization and effects on myogenic differentiation. PLoS One 5, e14447.

Valencia-Sanchez, M.I., De Ioannes, P., Wang, M., Truong, D.M., Lee, R., Armache, J.P., Boeke, J.D., and Armache, K.J. (2021). Regulation of the Dot1 histone H3K79 methyltransferase by histone H4K16 acetylation. Science 371.

Wang, Z.J., Rein, B., Zhong, P., Williams, J., Cao, Q., Yang, F., Zhang, F., Ma, K., and Yan, Z. (2021). Autism risk gene KMT5B deficiency in prefrontal cortex induces synaptic dysfunction and social deficits via alterations of DNA repair and gene transcription. Neuropsychopharmacology 46, 1617–1626.

Wilson, M.D., Benlekbir, S., Fradet-Turcotte, A., Sherker, A., Julien, J.P., McEwan, A., Noordermeer, S.M., Sicheri, F., Rubinstein, J.L., and Durocher, D. (2016). The structural basis of modified nucleosome recognition by 53BP1. Nature 536, 100–103.

Wu, H., Siarheyeva, A., Zeng, H., Lam, R., Dong, A., Wu, X.H., Li, Y., Schapira, M., Vedadi, M., and Min, J. (2013). Crystal structures of the human histone H4K20 methyltransferases SUV420H1 and SUV420H2. FEBS Lett 587, 3859–3868.

Xu, T.H., Liu, M., Zhou, X.E., Liang, G., Zhao, G., Xu, H.E., Melcher, K., and Jones, P.A. (2020). Structure of nucleosome-bound DNA methyltransferases DNMT3A and DNMT3B. Nature 586, 151–155.

Zheng, S.Q., Palovcak, E., Armache, J.P., Verba, K.A., Cheng, Y., and Agard, D.A. (2017). MotionCor2: anisotropic correction of beam-induced motion for improved cryo-electron microscopy. Nat Methods 14, 331–332.

Zhou, K., Gaullier, G., and Luger, K. (2019). Nucleosome structure and dynamics are coming of age. Nat Struct Mol Biol 26, 3–13.

